# An Asynchronous Production Line of Meiotic Prophase I in the Mouse Fetal Ovary

**DOI:** 10.64898/2026.01.14.699603

**Authors:** Chang Liu, Ziyi Jin, Gan Liu, Guofeng Feng, Jie Li, Yiwei Wu, Hao Jia, Lin Liu

**Author notes:** Corresponding author. (L.L.). These authors contributed equally to this work (Chang Liu, Ziyi Jin, Gan Liu).

## Abstract

The initiation of meiosis in the mammalian female germline has long been described as a synchronous event, occurring within a narrow developmental window. Here, we challenge this paradigm through a systematic, quantitative analysis of meiotic entry and progression in the mouse fetal ovary. Using dynamic expression profiling of key regulators Stra8, Sycp1, and Sycp3 alongside proliferation markers, we demonstrate that germ cells enter meiosis asynchronously and continuously between embryonic days E12.5 and E16.5. During this extended period, mitotic proliferation persists, indicating that germ cells are progressively recruited into the meiotic pathway rather than halting division simultaneously. Homologous chromosome synapsis, marked by Sycp1/Sycp3 co-localization, initiates at E14.5 and is completed prenatally by E18.5. Integrating these data into a continuous-time Markov chain model, we derive a “meiotic clock” that predicts fixed temporal intervals from meiotic entry to synapsis completion (∼72 hours) and to meiotic arrest (∼91 hours), regardless of entry time. This model provides a predictive framework for germ cell development and establishes that female meiotic initiation is a prolonged, asynchronous production line rather than a synchronized transition, reshaping our understanding of oogenesis and its regulatory logic.

**TEASER:** The initiation of female meiosis in mice is redefined as an asynchronous, continuous production line from E12.5 to E16.5, supported by a predictive “meiotic clock” model with fixed temporal intervals for synapsis and arrest.

## INTRODUCTION

Meiosis is the specialized cell division program that underlies gametogenesis and genetic diversity in sexually reproducing organisms. In mammals, the timing of meiotic initiation is sexually dimorphic: while spermatogenesis begins postnatally, oogenesis commences during fetal development(*1-4*). Female germ cells enter a state of arrest after completing the meiotic prophase I and only resume the process after puberty, rapidly completing the two meiotic divisions thereafter(*5*). Therefore, the transition from mitotic proliferation to meiotic prophase I in the female germline represents a pivotal developmental commitment, yet the precise dynamics and regulatory logic of this transition remain incompletely defined.

The prevailing model, supported by classical histology and marker expression studies, posits that germ cells in the fetal mouse ovary initiate meiosis between embryonic days 13.5 (E13.5) and 14.5 (E14.5), concomitant with a cessation of proliferation(*6-9*). This process is thought to occur in a synchronous, wave-like manner(*10, 11*). The model largely relies on the expression dynamics of key regulatory factors such as Stra8, a critical gatekeeper of meiotic entry(*12, 13*), and synaptonemal complex components like Sycp3, which labels chromosome axes during synapsis(*14*). Previous studies have established that Stra8, a central effector of the retinoic acid signaling pathway, plays an indispensable role in driving meiotic entry by regulating meiosis-related gene expression in both female and male germ cells(*6, 15-17*). A defining event of meiotic prophase I is the synapsis and recombination of homologous chromosomes, facilitated by the assembly of the synaptonemal complex (SC) starting at the zygotene stage(*7, 18*). Proteins such as Sycp1 and Sycp3 are essential for SC formation, serving as structural and functional markers of chromosome pairing and synapsis(*19-22*).

However, emerging evidence suggests a more complex, heterogeneous landscape of meiotic commitment, raising fundamental questions about the synchronicity, duration, and coordination of meiotic entry with mitotic activity. Critical gaps persist in our quantitative understanding of the protein-level dynamics of these regulators across the entire meiotic window, from initial commitment through synapsis to eventual prophase I arrest. Furthermore, the relationship between the lingering presence of proliferation markers and the progression of meiosis has been unclear, leaving open the possibility that germ cells may not embark on meiosis as a perfectly synchronized cohort.

Here, we perform a systematic, integrated analysis of protein expression, proliferation status, and single-cell transcriptomics to reconstruct the timeline of meiotic prophase I in the mouse fetal ovary. We challenge the canonical synchronous entry model by demonstrating that germ cells asynchronously and continuously enter meiosis over a four-day period from E12.5 to E16.5, while mitotic proliferation gradually declines. We define the precise window for homologous chromosome synapsis and integrate these data into a unified mathematical model—a “meiotic clock”—that reveals fixed, predictable intervals governing the progression from entry to synapsis to arrest. Our findings redefine the initiation of female meiosis as a prolonged production line, providing a quantitative framework for understanding oocyte development and its dysregulation.

## RESULTS

### Dynamic expression of the Stra8 and Sycp3 proteins in female fetal ovaries

To define the meiotic timeline more precisely, we collected female ICR mouse ovaries at stages E12.5, E13.5, E14.5, E16.5, E18.5, PD1 and PD5; performed serial sectioning and co-immunofluorescence for Stra8 for meiosis entry(*12, 23, 24*) and Sycp3(*25, 26*) for cohesion of synapsis homologs (Figures 1A, S1A); and quantified positive cells (Figure 1B, C). Stra8 expression was present from E12.5 onward, and the number of cells expressing Stra8 increased rapidly with increasing embryonic development, peaking at E13.5 and E14.5, after which the number of Stra8^+^ cells began to plummet, with a small number of cells still expressing Stra8 at E16.5, which was completely undetectable at E18.5 and thereafter (Figure 1B). These results imply that meiotic entry is not a transient, collectively synchronous entry process but rather a slow and long process starting at E12.5 and ending at E16.5 and that meiosis entry does not occur after birth. The number of Sycp3-positive cells continued to increase from E12.5 to E16.5 and then began to decrease but persisted until after birth (Figure 1C).

**Figure 1.**
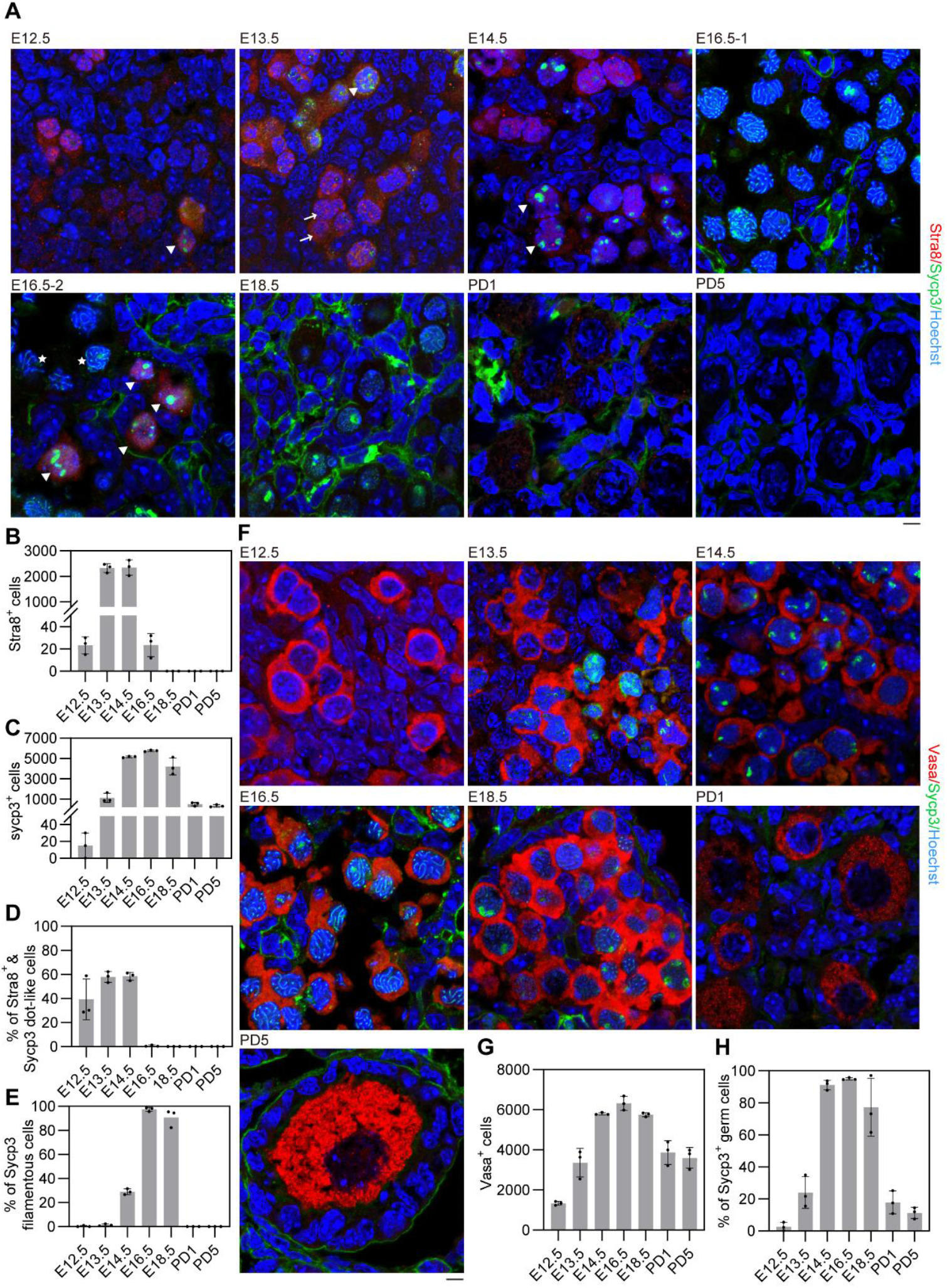
Dynamic expression of the Stra8, Sycp3 and Vasa proteins in ovaries at various embryonic stages. **(A)** Representative co-immunofluorescence images of Stra8 and Sycp3 in *in situ* sections from E12.5, E13.5, E14.5, E16.5, E18.5, PD1, and PD5 embryonic female gonads. Red, Stra8; green, Sycp3; blue, nuclei counterstained with Hoechst 33342. The arrows indicate Stra8 single-positive cells; the triangles indicate Stra8 and Sycp3 double-positive cells (Sycp3 dot-like distribution); the stars indicate Sycp3 single-positive cells (Sycp3 filamentous distribution); scale bars: 5 μm. **(B)** Statistical counts displaying the number of Stra8^+^ cells at various embryonic stages as indicated. **(C)** Statistical counts displaying the number of Sycp3^+^ cells at various embryonic stages as indicated. **(D)** Percentage (100%) of Stra8^+^ and Sycp3 dot-like cells, calculated as the number of Stra8^*+*^ and Sycp3 dot-like^*+*^ cells/sum of Stra8^*+*^ and Sycp3^*+*^ cells, including their co-expression. **(E)** Percentage (100%) of Stra8^−^ and Sycp3 filamentous cells, calculated as the number of Stra8^−^ and Sycp3 filamentous cells/Sum of Stra8^*+*^ and Sycp3^*+*^ cells, including their co-expression. **(F)** Representative co-immunofluorescence images of Vasa and Sycp3 in *in situ* sections from E12.5, E13.5, E14.5, E16.5, E18.5, PD1, and PD5 embryonic female gonads. Red, Vasa; green, Sycp3; blue, nuclei counterstained with Hoechst 33342. Scale bars: 5 μm. **(G)** Statistical counts showing the number of Vasa^+^ cells at various developmental stages as indicated. **(H)** Statistical counts showing the percentage of germ cells expressing Sycp3 at various developmental stages. **B, C, G, H**, Data are presented as the means ± SDs, n = 3 gonads in three independent experiments.

The data suggest a sequential expression pattern: Stra8 is expressed first (Figure 1A: arrows), followed by Sycp3, which initially appears as dot-like signals. At this stage, both Stra8 and Sycp3 coexist (Figure 1A: triangles). By E16.5, most cells reach pachytene, where Sycp3 exhibits a filamentous pattern and no longer co-expresses Stra8 (Figure 1A: stars). The percentage of Stra8^+^ Sycp3 dot-like cells gradually increased from E12.5 to E14.5, and then sharply decreased at E16.5. Conversely, the percentage of Sycp3 filamentous cells increased steadily, peaking at E16.5 (Figure 1D, E). Although most cells at E16.5 exhibit filamentous Sycp3 (pachytene), a subset remains in the Stra8^+^ Sycp3 dot-like state (Figure 1A, D), indicating that some germ cells have just initiated meiosis. This confirms asynchronous meiotic entry.

This sequential expression aligns with the role of Stra8 in meiotic initiation. Notably, prior to E13.5, the number of Sycp3^+^ cells was lower than that of Stra8^+^ cells, but this ratio reversed after E13.5 (Figure 1B, C). After E13.5, most cells respond to Stra8, followed by a surge of Sycp3^+^ cells by E14.5.

### A gradual increase in the occurrence of meiosis is accompanied by a gradual decrease in mitosis

Moreover, we performed co-immunofluorescence staining of Sycp3 with the germ cell marker Vasa (also known as Ddx4, a conserved marker commonly expressed in germ cells(*27-29*)) on consecutive ovarian sections to validate our quantitative results and examine meiotic progression (Figures 1F, S1B). All Sycp3-positive cells co-expressed Vasa. Quantitative analysis revealed that the number of germ cells was low at E12.5 (∼1,300 cells), progressively increased to a peak at E16.5 (>6,000 cells), and then declined postnatally to ∼3000-4000 (Figure 1G). The number of germ cells at E13.5 (∼3,500 cells) is consistent with that obtained via stereological estimation(*30, 31*), supporting our estimates. Notably, the decline in the proliferative rate of Vasa^+^ cells coincided with a decreased proportion of Stra8^+^/Sycp3^+^ dot-like cells entering meiosis (Figure 1D), suggesting that the number of germ cells capable of undergoing meiosis is reduced. The decreased number of germ cells after birth is also consistent with the findings of other studies(*32*).

We counted the proportion of Sycp3^+^ germ cells. At E12.5, only a very small number of germ cells entered the meiotic process, and then, the proportion of meiotic cells increased significantly. By E16.5, almost all existing germ cells had entered the meiotic process (Figure 1H). However, not all germ cells entered the meiotic process, and the proportion of cells that did not enter meiosis gradually decreased prior to E16.5. We speculate that some of these cells are very likely to be undergoing proliferation. Notably, the number of germ cells increased nearly three-fold from E12.5 to E16.5, and this dramatic proliferation prompted investigation of mitotic activity during meiotic initiation. Indeed, immunofluorescence of the proliferation markers Ki67 and Pcna(*33, 34*) with Vasa demonstrated active germ cell division during E12.5--E14.5 (Figures 2A, S2A, B). The number of Ki67^+^ Vasa^+^ cells peaked at E12.5, declined slightly at E13.5, and then decreased sharply by E14.5 (Figure 2B), whereas the number of Pcna^+^ Vasa^+^ cells peaked at E13.5 (Figure 2C). These patterns indicate sustained germ cell proliferation through E13.5, followed by marked reduction as meiosis advanced. Germ cell proliferation was completely inhibited by E16.5, which coincided with the completion of meiotic entry.

**Figure 2.**
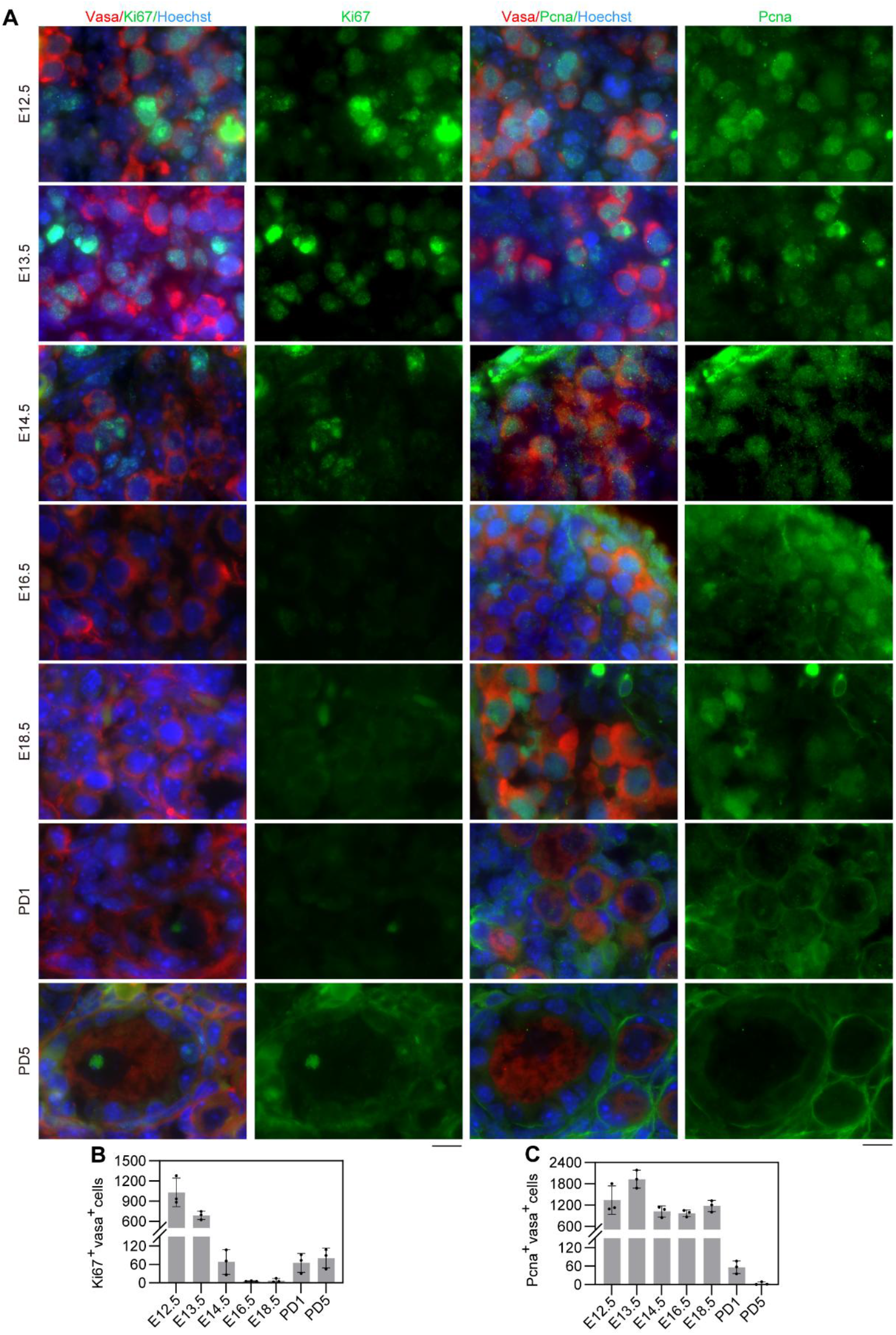
Dynamics of Ki67 and Pcna in germ cells of fetal ovaries. **(A)** Representative co-immunofluorescence images of Ki67 and Vasa or Pnca and Vasa in *in situ* sections from E12.5, E13.5, E14.5, E16.5, E18.5, PD1 and PD5 embryonic female gonads. Red, Vasa; green, Ki67 or Pcna; blue, nucleus counterstained with Hoechst 33342. Scale bars: 10 μm. **(B)** Statistical counts showing the number of Ki67^+^ Vasa^+^ cells at various stages as indicated. **(C)** Statistical counts showing the number of Pcna^+^ Vasa^+^ cells at various stages as indicated. **B, C**, Data are presented as the mean ± SD, n = 3 gonads in three independent experiments.

### Homologous chromosome synapsis and the frequency at various embryonic days

Synapsis is mediated by programmed interhomolog recombinational interactions in association with chromosome structural axes during prophase I and, ultimately, ensures the regular segregation of homologous chromosomes when they separate at the first meiotic division(*18, 35, 36*). To monitor chromosomal synapsis, we performed co-immunofluorescence of Sycp1 and Sycp3, which is based on a well-established method(*37, 38*), on ovarian sections from E12.5 to PD5 (Figures 3A, S3A). Distinct meiotic stages were identified through characteristic localization patterns *in situ* (Figure 3B), including leptotene (representative images at E12.5) exhibiting diffuse Sycp3 dot-like without Sycp1 expression, zygotene (E14.5) showing partial Sycp1 co-localization with Sycp3, which displayed both diffuse and short filamentous structures, pachytene (E16.5) demonstrating the complete co-localization of Sycp1 and Sycp3 as continuous filaments and diplotene (E18.5) maintaining Sycp3 filaments but showing Sycp1 disintegration, with residual Sycp1 localized at synapsed regions and Sycp3 exhibiting end-enriched fluorescence along chromosomes.

**Figure 3.**
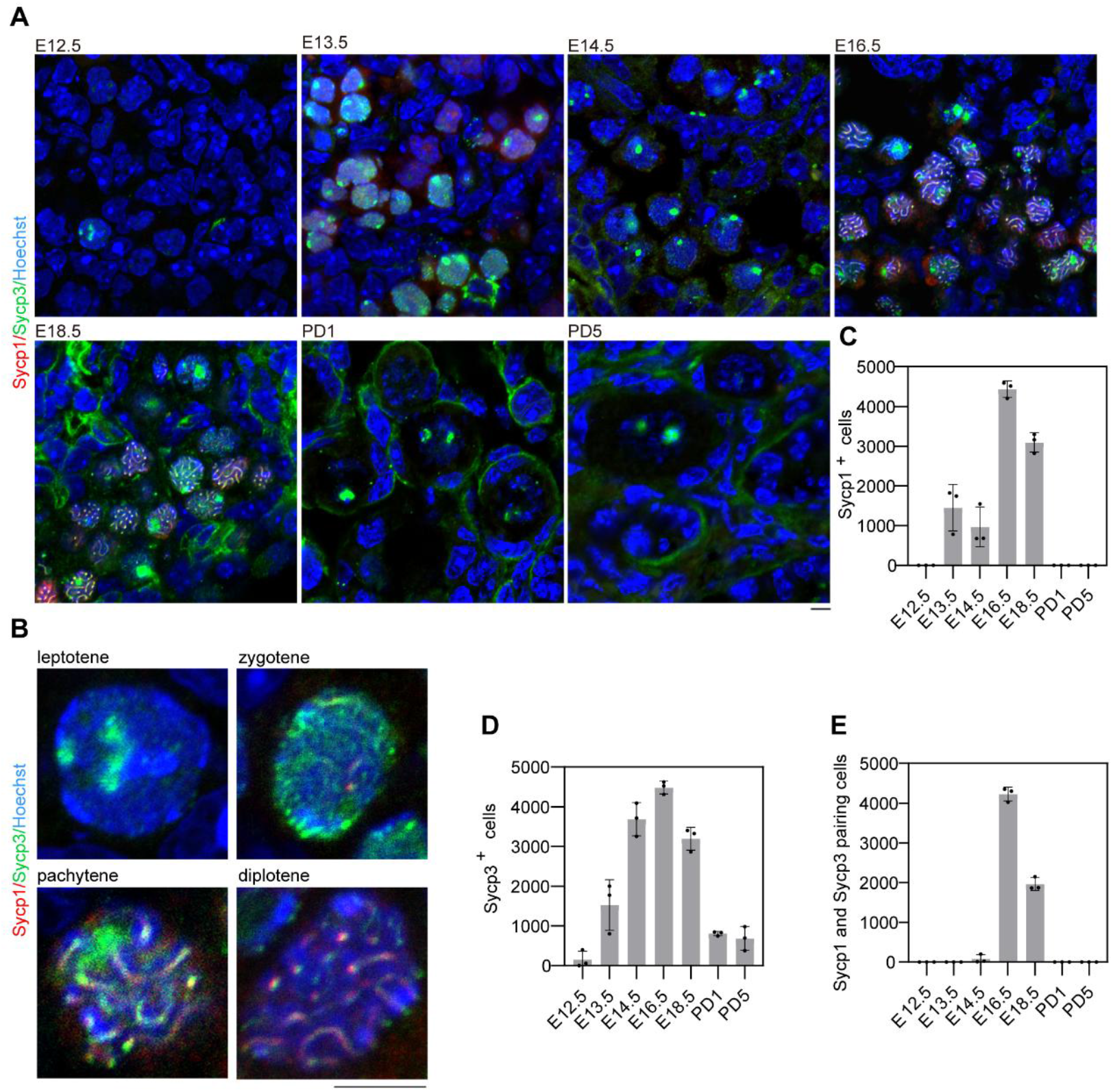
Dynamic expression of Sycp1 and Sycp3 and synapsis of Sycp1 and Sycp3 in ovaries. **(A)** Representative co-immunofluorescence images of Sycp1 and Sycp3 in *in situ* sections from E12.5, E13.5, E14.5, E16.5, E18.5, PD1, and PD5 embryonic female gonads. Red, Sycp1; green, Sycp3; blue, nuclei counterstained with Hoechst 33342. Scale bars: 5 μm. **(B)** Representative images of the morphology of meiocytes in the leptotene stage (from E12.5), zygotene stage (from E14.5), pachytene stage (from E16.5) and diplotene stage (from E18.5) observable in *in situ* samples. Red, Sycp1; green, Sycp3; blue, nuclei counterstained with Hoechst 33342. Scale bars: 5 μm. (**C-E**) Statistical counts showing the number of Sycp1^+^/Sycp3^+^/synapsed cells at various stages as indicated. Chromosome-synapsed cells were determined by intimate co-localization of Sycp1 and Sycp3. The data are presented as the means ± SDs; n = 3 gonads in three independent experiments.

To better characterize meiotic progression, we quantified Sycp1- and Sycp3-positive cells during various embryonic developmental stages. Sycp1 expression is initiated at E13.5, slightly later than Sycp3, initially displaying diffuse nuclear localization before being concentrated along chromosome axes by E14.5. This transition may be correlated with a modest decrease in the number of Sycp1^+^ cells at E14.5 (Figure 3C, D). Complete chromosomal synapsis, as indicated by filamentous Sycp1/Sycp3 co-localization, first appeared at E14.5 and then peaked at E16.5. By E18.5, while the number of synapsed Sycp1/Sycp3 was decreased, some cells with Sycp3 filaments showed only partial Sycp1 localization. Sycp1 became undetectable postnatally (PD1−PD5), whereas residual Sycp3 persisted. These temporal patterns demonstrate that the Sycp1 protein is expressed later than Sycp3 and that the Sycp1 protein ends earlier than Sycp3 does (Figure 3C-E). Sycp1 is critical for homologous synapsis.

### E14.5--E18.5 define the critical window for chromosome synapsis completion prior to meiotic arrest

To better characterize meiotic progression, we performed surface spread staining of E12.5 to PD5 ovaries (Figure 4A). This approach revealed distinct cytological features at each meiotic stage, preleptotene showing dot-like Sycp3 without Sycp1, leptotene with short Sycp3 filaments, zygotene displaying longer, discontinuous Sycp3 filaments with partial Sycp1 colocalization, pachytene with complete Sycp1/Sycp3 colocalization as thick filaments, diplotene exhibiting Sycp1 retention only at synapsed regions with visible separation, and at diakinesis, Sycp1 is completely degraded together with Sycp3 showing an intermittent distribution (Figure 4B).

**Figure 4.**
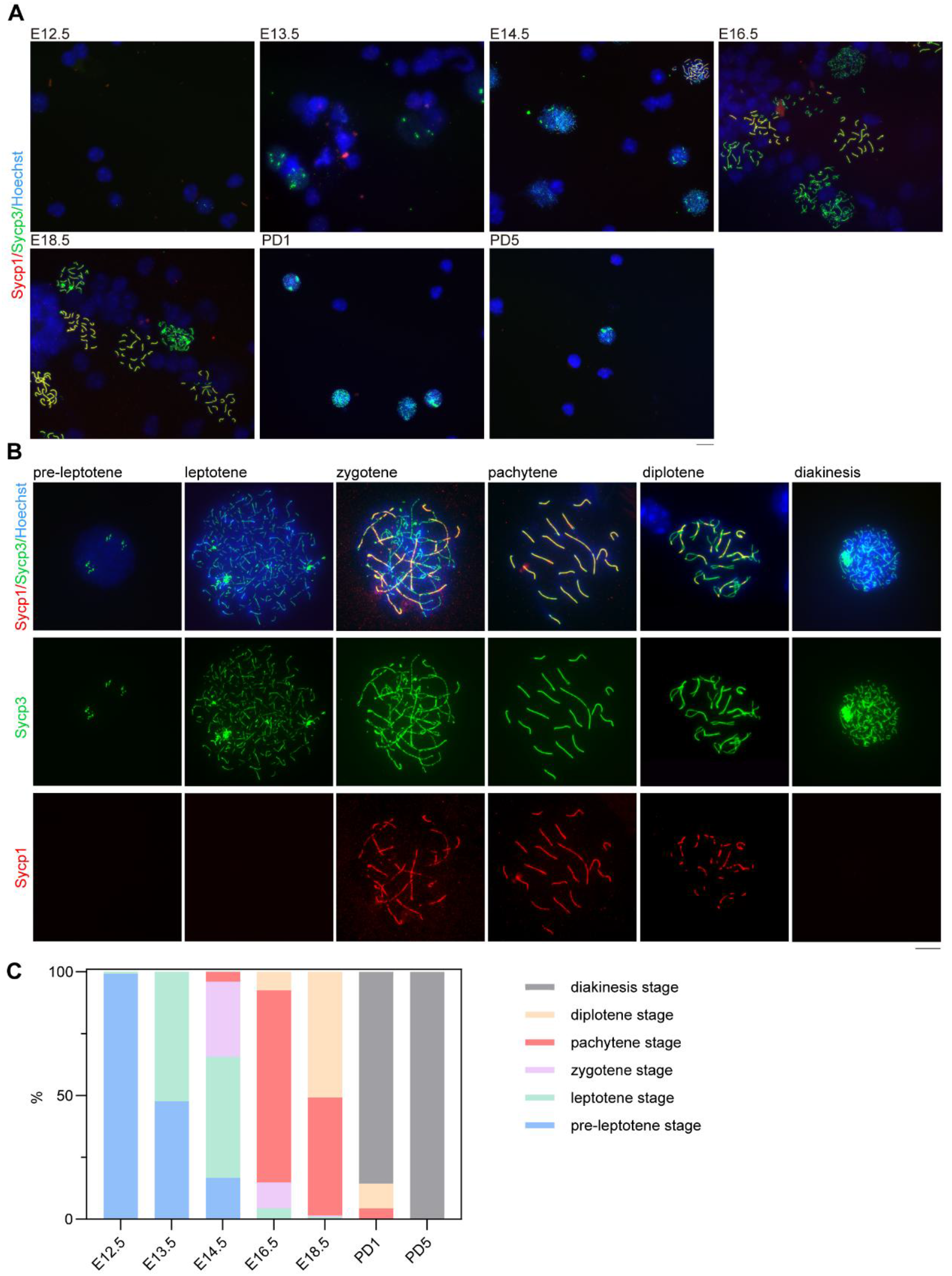
Estimation of the proportion of germ cells at different developmental stages via surface spread experiments. **(A)** Representative co-immunofluorescence images of surface-spread Sycp1 and Sycp3 from E12.5, E13.5, E14.5, E16.5, E18.5, PD1 and PD5 embryonic female gonads. Red, Sycp1; green, Sycp3; blue, nuclei counterstained with Hoechst 33342. Scale bars: 10 μm. **(B)** Representative images of germ cell typing based on Sycp1 and Sycp3 staining of female gonad surface spreads, where preleptotene cells are from E12.5, leptotene cells are from E13.5, zygotene cells are from E14.5, pachytene cells are from E16.5, and diplotene cells and diakinesis cells are from PD1. Red, Sycp1; green, Sycp3; blue, nuclei counterstained with Hoechst 33342. Scale bars: 10 μm. **(C)** Statistics on the proportion of germ cells at different developmental stages under surface spread. Twenty-five randomly selected visual fields were counted at each stage. The data are presented as the means, n = 3 repeats in three independent experiments.

Quantitative analysis of surface-spread meiocytes/germ cells revealed stage-specific progression through meiotic prophase I (Figure 4C). At E12.5, nearly all the germ cells remained in the preleptotene stage. The number of leptotene-stage cells markedly increased by E13.5, followed by the appearance of zygotene and rare pachytene cells at E14.5. Pachytene became the predominant stage at E16.5, although some leptotene and zygotene cells persisted, indicating ongoing meiotic entry. By E18.5, most germ cells had advanced to the pachytene or diplotene stages, indicating near-complete meiotic initiation. Postnatally, diakinesis-stage meiocytes predominated at PD1, with all cells arrested in diakinesis by PD5, marking the completion of prophase I progression and the establishment of meiotic arrest.

On the basis of the results of section staining and surface spread, we observed the morphology *in situ* and clearer structure of chromosomes at meiotic prophase I. The number of cells in the sections that displayed complete chromosome synapsis, as evidenced by Sycp1 and Sycp3, was mostly consistent with the proportion of cells at the pachytene stage, as determined in the chromosome spread experiment. However, the surface spread provides a clearer observation, indicating that synapsis of homologous chromosomes initiates at E14.5, continues to E18.5, and is completed at birth, followed by the onset of meiotic arrest.

### scRNA-seq analysis of germ cells during various developmental stages

To validate our findings, we analyzed published single-cell RNA-seq data from germ cells (*39, 40*). Clustering analysis revealed distinct meiotic stages: 0-leptotene, 1-diplotene, 2-pachytene, 3-zygotene, 4-diakinesis, and 5-premeiotic (Figure 5A, B). Temporal analysis revealed a progressive advancement of meiosis from E12.5 (predominantly at the leptotene stage) to E16.5 (primarily at the pachytene stage). By E18.5, most cells had reached the pachytene stage and eventually arrested at diakinesis after birth. In addition, a persistent population of pre-meiotic germ cells was observed throughout all stages examined.

**Figure 5.**
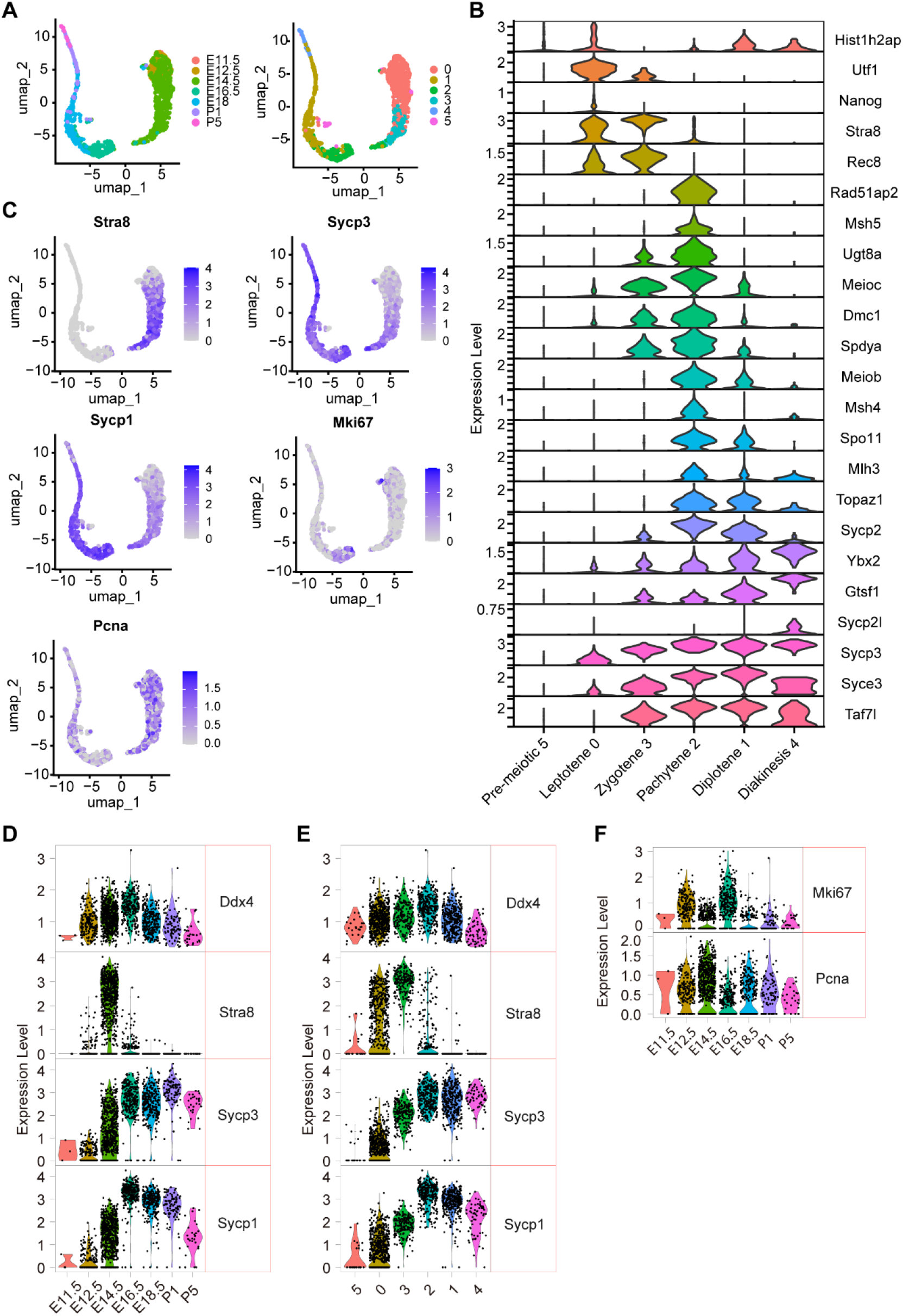
Analysis of 10x sc-RNA-seq data for expression levels of meiosis genes. **(A)** Visualization of clusters via umap. The cells are colored according to the embryonic time points from E11.5--P5 (left); the cells are colored according to the 6 identified cell groups (right). The clusters correspond to 0-leptotene; 1-diplotene; 2-pachytene; 3-zygotene; 4-diakinesis; and 5-premeiotic. **(B)** Multiviolin plot of selected meiosis-related gene expression during the six developmental stages. **(C)** Plots of the expression levels of *Stra8, Sycp3, Sycp1, Ki67* and *Pcna* along the developmental timeline; color intensity indicates the level of expression. **(D)** A multivariate plot showing the relative expression of genes (*Ddx4, Stra8*, Sycp3 and *Sycp1;* gene names at right) at different developmental time points (times at the bottom). **(E)** A multivariate plot showing the relative expression of genes (*Ddx4, Stra8*, Sycp3 and *Sycp1;* gene names at right) in cell clusters (clusters at bottom). **(F)** A multivariate plot showing the expression of *Ki67* and *Pcna* at different developmental time points (times at the bottom). All the 10x sc-RNA-seq data used here are from Niu et al.’s article(*39*).

The transcriptional profile revealed that *Stra8* is predominantly expressed in the early part of the prophase I event, whereas the components of the synaptonemal complex are expressed throughout the E12.5−PD5 period. During this period, the proliferation markers *Ki67* and *Pcna* were also expressed (Figures 5C, S4A). The specific expression trends indicate that *Ddx4* is expressed throughout all periods. *Stra8* mRNA levels peak at E13.5--E14.5. The mRNA expression levels of the synaptonemal complex components continued to increase before E16.5 (Figures 5D, S4B). *Ddx4* has comparable expression levels throughout all developmental stages, while *Stra8* shows maximal expression during the leptotene–zygotene transition. *Sycp3* and *Sycp1* mRNA levels increased from leptotene through later stages and peaked at the pachytene stage (Figure 5E). Moreover, proliferation markers such as *Ki67* and *Pcna* are expressed at relatively high levels from E12.5 to E16.5 (Figures 5F, S4C), which is consistent with the expansion of germ cells during this period. This comprehensive single-cell analysis not only validated our histological findings but also provided molecular resolution of meiotic progression and concurrent proliferation events.

### Parametric modeling and prediction of marker timing and stage composition

We quantified immunostaining trajectories for germ-cell markers across seven time points from embryonic day E12.5 to postnatal day PD5. For each panel, the proportion of marker-positive cells was computed using the number of Vasa+ cells as the denominator, and replicate panels at each time point were pooled by denominator-weighted aggregation. The trajectories revealed a sequential pattern from meiotic entry to synapsis and axial compaction. Stra8 exhibited an early, narrow window with t10↑– t90↑≈E12.58–E13.40, peaking at E13.50 (peak≈0.69), and the FWHM≈1.81 days. Sycp3 (filamentous) peaked nearly synchronously with Sycp1 at E16.50 (peak≈0.70) but with a shorter FWHM≈3.68 days; in contrast, Sycp3 (dot-like) dominated earlier (peak E14.50, peak≈0.59). Ki67 was highest at E12.5 (≈0.77) and rapidly declined; Pcna was close to 1.00 at the earliest time point and was used primarily as an auxiliary display (Figure S5 A-B, Table S1). To quantify precedence, we compared rising half-times across markers, yielding lags Δt50↑that capture “who leads and by how much.” We subsequently integrated replicate data across time points via a denominator-weighted pooling method (numerator and denominator summed at the time level) to quantify the meiotic stage composition at seven time points from E12.5--PD5, thereby obtaining both the observed stacked composition and the corresponding model-predicted results (Figure S5C−D, Table S2−4). The fitted trajectories delineate a sequential relay from leptotene (L) to diplotene (D). Z exceeds L at 14.70, P exceeds Z at 15.55, and D exceeds P at 18.46. These relay crossings, together with per-stage windows, quantify the order and span of progression. Model-based prediction enables interpolation/extrapolation at unmeasured times.

### An interpretable meiotic clock: CTMC-based stage dynamics, molecular alignment, and steep Dip-to-Dia hand-off

At the aforementioned time points, the interval solutions obtained via the piecewise CTMC are highly consistent with the observed stage compositions (Figure 6A, solid lines represent the model, and scatter points represent observations). The model is dominant in the *q*_*LZ*_ segment during E13.5–E14.5, reproducing the narrow peak of *π* _*Z*_ (*t*) near E14.5. Subsequently, *π* _*P*_ (*t*) begins to rise rapidly around E15, forms a single peak during E16.5–E18.5, and then decreases after E19, reflecting a piecewise transition of *q*_*ZP*_ rather than a global smooth transition. For *π* _*D*_ (*t*), the interval solution shows an S-shaped accumulation: it crosses 0.5 during E18.5–PD1 and approaches 1.0 during PD1–PD5, indicating that the contribution of *q*_*PD*_ is concentrated mainly in the later stages. Overall, this piecewise-constant model accurately recaptures the key dynamics of all three stages (Z, P, D) with minimal parameters, establishing a unified “stage clock” for precise temporal quantification. Within this unified stage clock framework, molecular expression profiles were accurately predicted (Figure 6B, Table S5). The increasing half-width of Stra8 ( *t*50_up_ = 13.41, FWHM 1.33 days) indicates that a narrow expression window occurred in the early stage. Sycp1 displayed a broad expression profile ( *t*50_up_ = 13.46, FWHM: 5.41 days), featuring both a sharp early peak (∼13.5–14.5 days) and a main phase that mirrored Z→P progression while gradually decaying, reflecting its wider, more persistent window. *Sycp3_fil* shows a later-onset, intermediate-duration expression window ( *t*50_up_ = 15.40, FWHM 3.67 days), and *Sycp1*’s main peak aligns with *Sycp3_fil*. The overlapping region of Sycp1 and Sycp3 aligns with the mid-phase peak morphology of P, further validating the chromosomal architecture characteristics of the pachytene stage. Based on the rise half-duration, Sycp3_fil lagged behind Stra8 and Sycp1 by approximately 1.99 and 1.94 days, respectively. This temporal hierarchy aligns with the fixed-interval logic of “entry (Stra8/early Sycp1) → synapsis and advancement of pachytene (main Sycp1) → axis condensation (Sycp3_fil) → arrest (D)”. Thus, the ‘entry → pairing → arrest’ rhythm is not only reflected in the CTMC framework of morphological stages but also encoded in the temporal trajectories of molecular expression.

**Figure 6.**
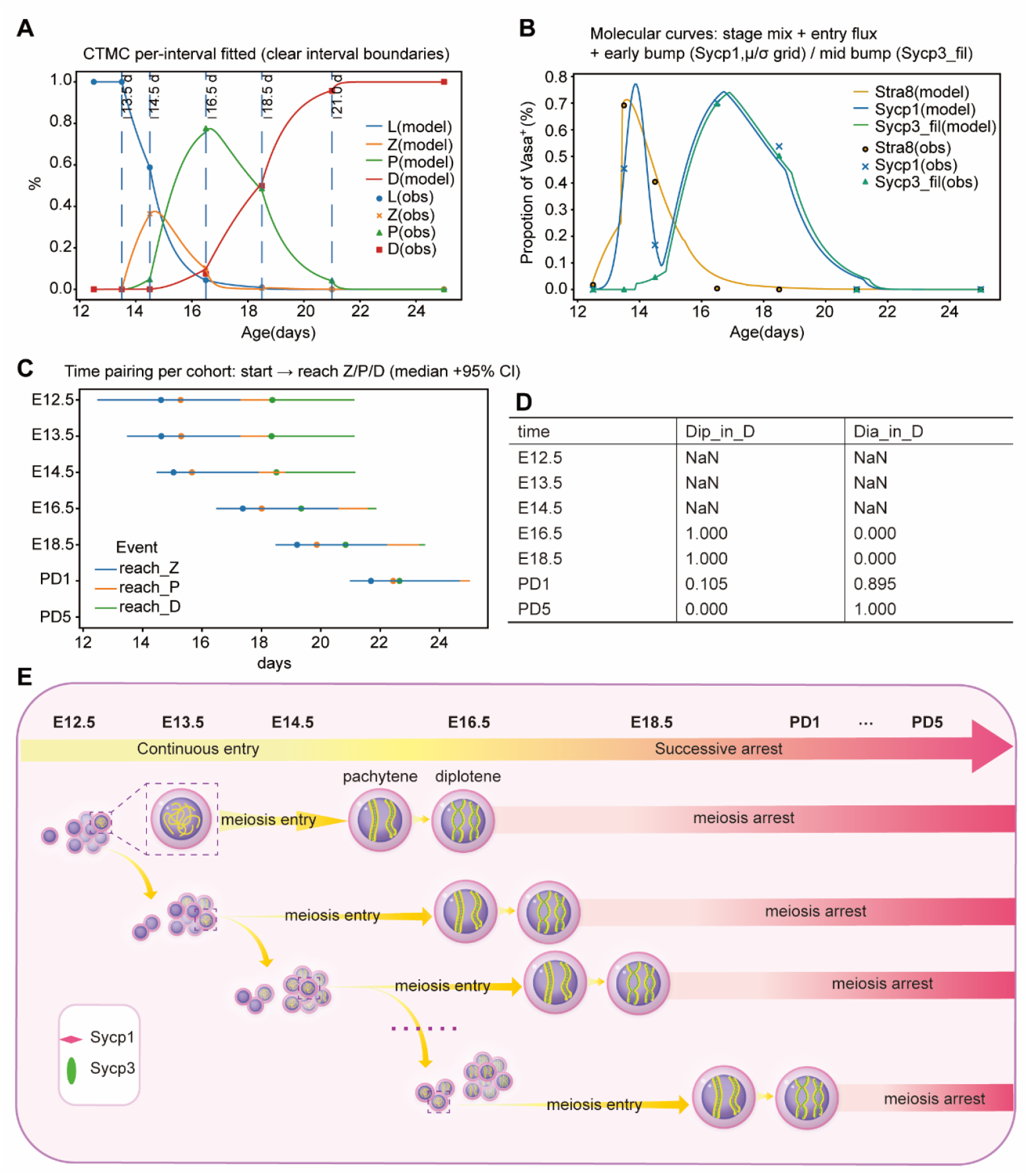
Modeling meiotic prophase I in female mice via CTMCs. **(A)** Segmented CTMC fitting on a unified phase clock (L→Z→P→D). The solid line represents the model-predicted stage composition **π**(*t*) = [*π* _*L*_,*π* _*Z*_,*π* _*P*_,*π* _*D*_ ], while the scatter points indicate the observed stage composition at the corresponding time points; the vertical dashed lines denote the boundaries between adjacent time intervals. The model assumes only forward transitions, with a constant rate matrix within each time interval [*t*_*k*_, *t*_*k*+1_ ), and ensures the continuous connection of segments at the boundaries. **(B)** Molecular–morphological alignment: Predicting molecular proportions via ‘stage composition + entry flux + local terms’ on a unified stage clock. The solid line represents the predicted molecular proportions 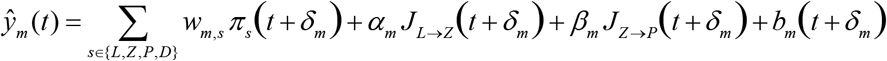 Stra8, Sycp1, Sycp3_fil), whereas the scatter points denote the observed molecular proportions *y*_*m*_ (*t* )(all normalized by *Vasa*^+^ ). A Gaussian local term with constrained amplitude was applied to Sycp1 to account for the early small peak at 13.5--14.5 days; the sharp condensation peak of Sycp3_fil at approximately days 16--18 was captured in the model by a narrow Gaussian local term centered in the middle stage; all three curves align with morphological progression on the unified stage clock. **(C)** Arrival time on the basis of cumulative flux: time scale for each cohort from start to Z/P/D. Each row represents a cohort of cells starting at a specific postnatal day; the horizontal lines indicate the relative arrival time ranges (median with 95% bootstrap confidence intervals) from the cohort start to reach the Z, P, and D stages, with the “×” marker denoting the median. The relative arrival times remain stable across different starting points, whereas the absolute arrival times shift with the starting point, revealing a fixed time interval scale for the “entry → pairing → arrest” sequence. **(D)** Internal composition of Stage D. The Dip–Dia ratio at various time points when *N*_*D*_ = 0 or not measured is defined as NaN (E12.5–E14.5). **(E)** Schematic diagram of germ cells continuously entering meiosis.

For the unified stage clock, which defines the arrival times for each stage via the cumulative flux *C*_*S*_ (*t*; *a*_start_ ), we obtain relatively stable time intervals across cohorts. The median values summarized by bootstrapping show that, measured from the cohort start *a*_start_, the relative arrival time to reach Z is 2.84 days (95% CI: 2.62–3.48 days, i.e., 62.9–83.5 hours), that to reach P is 3.01 days (2.79–3.70 days, 67.0–88.8 hours), and that to reach D is 3.79 days (3.54–4.49 days, 85.0–107.8 hours). This means that the three “fixed intervals” can be directly derived from the data: from “entry into meiosis (starting point L) → completion of pairing (P)” is approximately 3.01 days ≈ 72.2 hours; from “entry into meiosis (starting point L) → arrest (D)” is approximately 3.79 days ≈ 91.0 hours. Figure 6C further shows that the horizontal segments for different starting cohorts shift almost in parallel on the same clock: the relative arrival times remain stable across different starting points, whereas the absolute arrival times 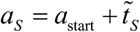 shift linearly with the starting point. Applying the aforementioned “fixed intervals” to specific starting points directly yields absolute predictions for “entry on day X → when pairing/arrest occurs”, which are consistent with the markers in the figure: if entry into meiosis occurs at E12.5, pairing is predicted to complete at E15.5 (≈E12.5 + 3.01) and arrest at E16.3 (≈E12.5 + 3.79); if entry occurs at E13.5, pairing is predicted to complete at E16.5 and arrest to begin at E17.3.

The normalized Dip_in_D and Dia_in_D series show a rapid transition from Dip dominance to Dia dominance between E18.5 and PD1 (Figure 6D). This shift aligns precisely with the predicted arrest times (E18.3–E20.3) for cohorts entering meiosis at E14.5–E16.5. Earlier and later cohorts arrest in pre-or post-transition stages, respectively, demonstrating that population-level Dip→Dia remodeling is a superposition of multiple cohorts undergoing arrest with a fixed ∼91-hour delay from their entry rather than a synchronized event.

In summary, through unified temporal modeling, we simulated the complete process of female meiotic progression in mice during embryonic development, from initial entry through pairing and synapsis to eventual arrest (Figure 6E).

## DISCUSSION

The initiation of female mammalian meiosis represents a fundamental divergence from the male germline program, commencing during fetal development and progressing through meiotic prophase I stages (leptotene, zygotene, pachytene, diplotene and diakinesis) before arresting until postnatal sexual maturation (*41, 42*). While the general sequence of meiotic events is well characterized(*43*), critical gaps persist in defining precise molecular criteria for meiotic entry. Prevailing models often presume synchronous initiation of meiosis across the oogonial population(*44*). Our integrated analysis of Stra8, Sycp1, Sycp3, and proliferation markers revealed that meiotic entry in mouse ovarian germ cells occurs as a temporally extended process spanning E12.5 to E16.5. Beginning at E12.5, germ cell populations diverge into distinct developmental trajectories, with a subset initiating meiosis and other uncommitted germ cells maintaining mitotic proliferation. This mixed population state persists through subsequent developmental stages, with proliferating cells continuously facing fate decisions between continued mitosis or meiotic commitment. This situation persists throughout the four-day developmental window, with mitotic activity only ceasing completely upon completion of meiotic entry by E16.5, at which point all germ cells have entered meiosis, mitotic proliferation ceases, and the total germ cell number stabilizes.

These results indicate that the initiation of meiosis is achieved through a gradual process rather than a simultaneous transformation process. This paradigm may support the model proposing wave-like meiotic initiation. Menke and Bullejos et al. previously identified an anterior-to-posterior differentiation wave through Stra8 expression patterns from E12.5 to E16.5(*10*). The meiotic commitment constitutes an extended temporal sequence rather than an instantaneous population-wide event.

Furthermore, we systematically quantified the expression patterns of the Sycp1 and Sycp3 proteins, which are key for the synapsis of homologous chromosomes. The sustained expression of Sycp3 during meiosis prophase I is consistent with the flow cytometry results(*45*), thus confirming the accuracy of our statistics. We show that homologous chromosome synapsis, characterized by Sycp1 and Sycp3 co-localization, initiates at E14.5 and persists until E18.5. However, the number of synapsis events detected *in situ* at E14.5 via this method was relatively low. In contrast, surface spread chromosome preparations revealed a significantly greater proportion of germ cells exhibiting homologous chromosome synapsis (in the zygotene and pachytene stages) at E14.5. This discrepancy is likely due to the lower Sycp1 expression during zygotene, making synapsed cells less detectable and potentially overlooked in sectioned tissue analysis. Collectively, these findings demonstrate that a substantial proportion of germ cells undergo homologous chromosome synapsis as early as E14.5. Overall, we established a reference timeline for the occurrence of homologous chromosome synapsis during E14.5–E18.5. The absence of Sycp1 at PD1 and PD5 clearly indicates that homologous chromosomes cease pairing and synapsis postnatally.

Through calculation, we obtained an extrapolatable mathematical model of protein expression and discovered that meiotic progression follows an intrinsic and quantifiable timing program. Using a unified stage clock, we determined the key fixed intervals from meiotic entry to pairing completion (∼72 hours) and to meiotic arrest (∼91 hours). These intervals remain consistent across cell populations with different starting points, thereby transforming the complete sequence—from the end of proliferation through meiotic entry, pairing, and arrest—into a predictable temporal scale. This model is mutually corroborated by observed molecular expression windows and internal stage-D compositional dynamics, and its predictive ability provides a reliable framework for further experimental validation and the design of temporal interventions.

Our findings collectively redefine the paradigm of meiotic initiation in mammalian oogenesis by demonstrating that germ cells undergo a progressive, asynchronous recruitment into the meiosis production line rather than a synchronous “all-or-none” transition. However, the molecular mechanisms driving the production line of meiosis I remain to be fully elucidated. In this study, all experimental data were derived exclusively from ICR mice; the precise timing of meiotic entry may vary across different mouse genetic backgrounds. Future technological advances could enable more accurate recording of comprehensive gonad-wide data.

## MATERIALS AND METHODS

### Mice

ICR mice were purchased from Beijing Weitong Lihua Company and were confirmed to have no specific pathogens. The mice used in this study followed the relevant regulations of the Experimental Animal Center and the Animal Ethics Committee of Nankai University, and all experiments were approved by the Animal Ethics Committee. All mice were maintained in the specific pathogen-free animal facility of the Experimental Animal Center, Nankai University, with a humidity of 35%±4%, a stable temperature of 24±1°C, and a 12/12 h light/dark cycle (8:00–20:00 light). Male mice were housed separately in each cage, while female mice were housed 5 per cage.

### Acquisition and separation of gonads at various developmental stages

Six-week-old female mice were subsequently mated with eight-week-old male mice (2:1) overnight. During the periods of 08:00 and 09:00 on the second day, the presence of vaginal plugs was checked, which was recorded as E0.5(*46*). All animals were humanely euthanized by cervical dislocation for tissue collection. Death was further assessed by confirmation of rigor mortis according to AVMA Guidelines for the Euthanasia of Animals. All efforts were made to minimize the number of animals used and their suffering.

After they reached the corresponding stages (E12.5, E13.5, E14.5, E16.5, and E18.5), the pregnant mice were euthanized, and the embryos at the corresponding stages were collected in PBS. For newborn mice (PD1, PD5), direct euthanasia was performed. The embryonic urogenital complexes were carefully removed under a stereomicroscope. The embryonic gonads were separated from adjacent mesonephros via syringe needles. We attempted to ensure that no other tissues were attached around the ovaries. After two rounds of washing with PBS, the intact gonads were collected in 1.5 ml EP tubes.

### Continuous frozen sections of gonads

The ovarian tissue was fixed as described previously(*12*). The female gonads were fixed in 4% PFA (paraformaldehyde) at 4°C overnight, the 4% PFA was removed, and the mixture was then replaced with 30% sucrose for dehydration for 2 hours. Once the tissue has settled, it should be embedded in optimal cutting temperature (OCT) ice. The sample can then be stored at -80 degrees Celsius or directly frozen.

In accordance with the method for counting follicles, continuous slicing should be performed at a thickness of 5 μm until the entire gonad is sectioned. Each slice is ensured to be complete and smooth to ensure credibility. One out of every five sheets should be taken for counting (*47, 48*). The gonads from each period were taken from different individuals, and the process was repeated three times.

### Immunofluorescence microscopy of frozen sections

Immunofluorescence microscopy of ovaries was performed mainly according to a previously described procedure(*49*). Specifically, the slides were incubated at room temperature for 30 min, washed in PBS for 10 min, fixed with precooled acetone for 15 min, dried in a fume hood for 10 min, permeabilized for 30 min in PBS containing 0.1% Triton X-100, washed twice with PBS, blocked with 3% BSA in PBS at room temperature in a humid box for at least 2 h, and then incubated with primary antibodies against Vasa (ab13840, Abcam, 1:200), Vasa (ab27591, Abcam, 1:200), Stra8 (ab49602, Abcam, 1:200), Sycp3 (ab97672, Abcam, 1:200), Sycp1 (ab303520, Abcam, 1:200), Pcna (ab29, Abcam, 1:200) or Ki67 (ab16667, Abcam, 1:200) overnight at 4°C. The next day, the humid box was removed from the 4°C refrigerator to warm to room temperature. The samples were washed three times with PBS (each time for 15 minutes) and incubated at room temperature in the dark for 2 hours with the appropriate fluorescence-conjugated secondary antibodies: donkey anti-rabbit IgG (H+L) Alexa Fluor 594 antibody (Thermo Scientific, A-21207, 1:200), donkey anti-mouse IgG (H+L) Alexa Fluor 488 antibody (Thermo Scientific, A-21202, 1:200) or donkey anti-mouse IgG (H+L) Alexa Fluor 594 antibody (Thermo Scientific, A-21203, 1:200), and donkey anti-rabbit IgG (H+L) Alexa Fluor 488 antibody (Thermo Scientific, A-21206, 1:200). The samples were washed three times with PBS (each time for 15 minutes) and counterstained with Hoechst 33342 (H1398, Molecular Probes) at a concentration of 0.5 mg/mL in Vectashield (Vector Labs) mounting medium. The fluorescence was detected and imaged via an Axio Imager Z2 fluorescence microscope (Zeiss) or a confocal microscope (Zeiss LSM710).

### Surface Spread

The surface of female ovaries was spread, and the samples were stained for synaptonemal complexes, mainly according to a previously described procedure (*50, 51*). The specific steps were as follows: the harvested tissues were digested in 0.05% TE at 37°C for 7 minutes, the digestion was terminated, centrifugation was performed, the supernatant was gently removed, and then an appropriate 100 mM sucrose solution was added to resuspend the precipitate for 3–5 minutes to cause the cells to burst. Clean slides were prepared, a border was drawn on the slide with a hydrophobic pen, and first, 1% PFA was added, followed immediately by gently tilting the slide with the resuspended sucrose solution to spread the solution evenly and placing it in a humid box at room temperature for 3 hours. After 3 hours, the cover was opened, and the samples were allowed to air dry slowly. The slides were washed with 0.4% photo flow for 2 minutes and air dried. Then, the samples were subjected to PBS containing 0.1% Triton X-100 for 10 minutes, blocked with 3% ADB solution for 2 hours at room temperature, and incubated with primary antibodies in blocking solution against Sycp3 (ab97672, Abcam, 1:200) and Sycp1 (ab303520, Abcam, 1:200) overnight at 4°C. The next day, the humid box was removed from the 4°C refrigerator to warm to room temperature. The samples were washed in PBS 3 times and incubated at room temperature in the dark for two hours with the appropriate fluorescence-conjugated secondary antibodies: donkey anti-rabbit IgG (H+L) Alexa Fluor 594 antibody (Thermo Scientific, A-21207, 1:200), donkey anti-mouse IgG (H+L) Alexa Fluor 488 antibody (Thermo Scientific, A-21202, 1:200), donkey anti-mouse IgG (H+L) Alexa Fluor 594 antibody (Thermo Scientific, A-21203, 1:200), and donkey anti-rabbit IgG (H+L) Alexa Fluor 488 antibody (Thermo Scientific, A-21206, 1:200). The samples were washed three times in PBS and counterstained with 0.5 mg/mL Hoechst 33342 (H1398, Molecular Probes) in Vectashield (Vector Labs) mounting medium. The fluorescence was detected and imaged via an Axio Imager Z2 fluorescence microscope (Zeiss) or a confocal microscope (Zeiss LSM710).

### 10 × Genomics computational analysis

The raw count matrices were imported into R for further processing. R Studio (https://www.rstudio.com/) was used to execute R scripts for conducting hierarchical clustering and PCA. To identify distinct cell populations of germ cells at different timepoints, cell clustering was performed via the R software package Seurat 5.0(*52*). The count matrix was initially normalized by library size and log transformed by Seurat. Transcriptomes with <1000 expressed genes were discarded, whereas those with mitochondrial genes accounting for >5% of the reads were defined as low-quality cells and were filtered out. Different timepoints of germ cells were integrated via the “IntegratedData” function of Seurat according to the instructions. Uniform manifold approximation (U-MAP) was employed for visualization and clustering. The “FindAllMarkers” function was utilized to identify cell-type marker genes that are specific across conditions. The “VlnPlot” function was used to plot the expression of proteins associated with different stages of meiosis, and the cell groups were categorized on the basis of the expression of proteins at various stages. All of the 10x sc-RNA-seq data used in this study are from Niu et al. and Shen et al. (*39, 40*).

### Data and normalization

At each time point *t* ∈{*E*12.5, …, *PD*5}, we recorded, per panel *i*, the counts 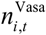 and 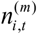 for markers *m* ∈{Stra8,Sycp1,Sycp3_dot_,Sycp3_fil_, Ki67, PCNA}. The panel-level proportion was 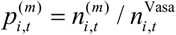. The pooled proportion at time *t* was the denominator-weighted estimator

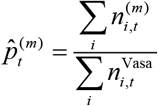

with Wilson 95% confidence intervals reported for pointwise uncertainty. Missing values were omitted from pooling.

Feature definitions. Let *f*^(*m*)^ (*t*) denote the (smoothed or fitted) trajectory. We defined:

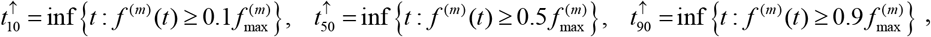

Where 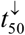 is defined symmetrically on the falling segment and 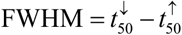. Marker precedence was summarized by 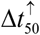 between the markers.

### Parametric two-logistic model

To support interpolation/extrapolation and one-to-one mapping to window metrics, we model “rise–then–fall” trajectories via a product of logistic functions:

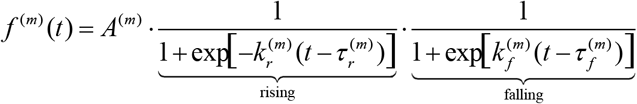

Where *A*^(*m*)^ is the amplitude, 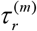 and 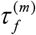 approximate 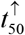 and t 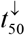, and 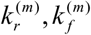 controls the steepness of the rising and falling limbs. Model parameters map one-to-one to window metrics, enabling direct estimation of 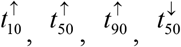, and FWHM as well as 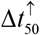 . The fitted curves agreed well with the observations for Stra8, Sycp1 and Sycp3 (filamentous), and the parameter estimates are summarized in Table S3.

### Counts, pooling, and QC

At time t, replicate spreads contributed counts of cells in stages *k* ∈{*L, Z, P, D*} with total *N*_t_. Time-level composition was estimated by pooling numerators and denominators across replicates under prespecified QC: removing failed stain/imaging fields, dropping replicates with very small totals (e.g., <20), and excluding records with counting inconsistencies; missing values were ignored. Postnatal days were mapped onto the same numeric axis as embryonic days.

### Multinomial logit with natural cubic splines

Let *π* _*k*_ (*t*) be the true proportion in stage k at time t, where 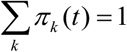. Using D as the baseline class for identifiability, we model the log odds of *L, Z, P* relative to D as smooth functions of time via a natural cubic spline basis 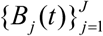.

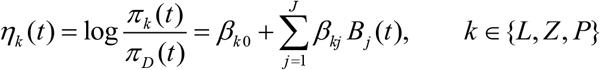

where *t* denotes developmental time (days); *k* ∈{*L, Z, P*} indexes meiotic stages— leptotene (L), zygotene (Z), and pachytene (P); *π* _*k*_ (*t*) is the true proportion of stage *k* at time *t* with; *D* is the baseline stage for identifiability; *η*_*k*_ (*t*) = log{*π*_*k*_ (*t*) / *π*_*D*_ (*t*)} is the log-odds for stage *k* relative to *D* ; 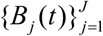 are natural cubic spline basis functions of time; *J* is the number of spline basis functions; and *β*_*k* 0_ and *β*_*kj*_ are regression coefficients. The implied softmax reconstruction gives:

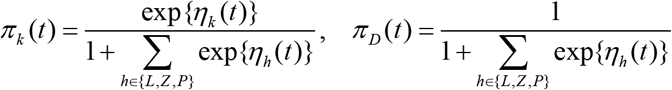

where *h* ∈{*L, Z, P*} indexes nonbaseline stages in the Softmax denominator and where exp{·} denotes the natural exponential function.

This specification yields smooth, fully probabilistic trajectories {*π* _*k*_ (*t*)} over time.

### Crossing times and reporting rules

The crossing times between stages *a,b* were defined as solutions to *π* _*a*_ (*t*) = *π*_*b*_ (*t*) ; to avoid spurious low-tail intersections, we reported only crossings where both *π* _*a*_ (*t*),*π*_*b*_ (*t*) ≥ 0.05. We summarized three relay anchors: *Z* > *L, P* > *Z*, and *D* > *P* .

### Data integration and notation

At discrete time points

𝒯 = {12.5,13.5,14.5,16.5,18.5,21.0,25.0} (days)

where 𝒯 denotes the sampling age in days, commonly represented as E12.5…PD5 in the text.

Stages, Stage Counting, and Stage Composition

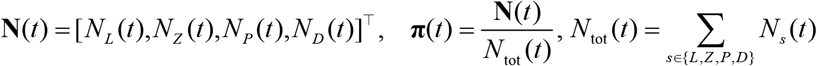

Where *N*_*s*_ (*t* )denotes the cell count of *s* ∈{*L, Z, P, D*} at time point *t* ; **π**(*t*) = [*π*_*L*_ (*t*),*π*_*Z*_ (*t*),*π*_*P*_ (*t*),*π*_*D*_ (*t*)]^⊤^ represents the stage composition; and **π**(*t*) denotes the stage composition vector (with components *π* _*s*_ (*t*) ∈[0,1] and 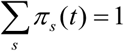 ). The scatter plot in Figure 6A corresponds to **π**(*t*).

D Internal Subtyping

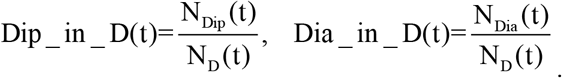

where *N*_Dip_ (*t*), *N*_Dia_ (*t*) represents the count of the two subtypes within D; when *N*_*D*_ (*t* ) = 0 or is undefined, the above expression is defined as NaN.

Ratio

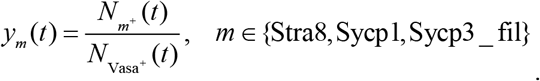

where 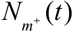 denotes the number of *m* positive cells; 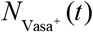 denotes the total number of *Vasa*^+^ ells; and *y*_*m*_ (*t*) ∈[0,1].

### Segmented CTMC Stage backbone

Structure and assumptions

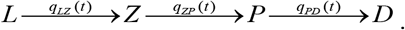

where q *q*_*XY*_ (*t* )is the instantaneous transition rate of *X* → *Y* (unit: *day*^−1^ ). Segmented constant rate

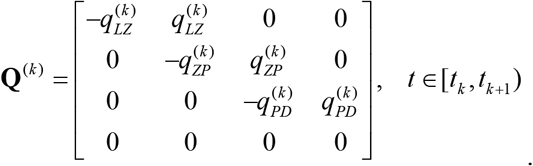

where the rate constant for the *k* segment [*t*_*k*_, *t*_*k* +1_ ) is *q*^(*k* )^ ≥ 0 and where is the 4×4 rate matrix.

Interval solution composed of stages

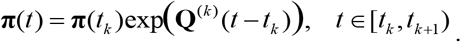

where exp(·) denotes the matrix exponent; **π**(*t*_*k*_ ) represents the initial value for this segment; and the segments are continuously connected at point *t*_*k* +1_ .

Parameter estimation within segments

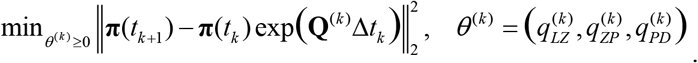

Where Δ*t*_*k*_ = *t*_*k* +1_ − *t*_*k*_ ;‖·‖× _2_ denotes the L2-norm; and *θ*_(*k* )_ is bounded within [0,5]. Enter flux

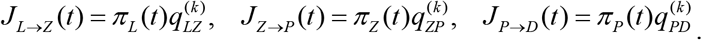

where *J*_*X* →*Y*_ (*t*) is the proportion of the population entering stage *Y* per unit time. Cumulative Flux (Relative Starting Point **a** )

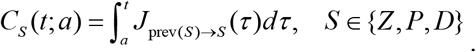

where prev(*Z* ) = *L*, prev(*P*) = *Z*, prev(*D*) = *P* .

### Flux aperture ‘arrival time’

Relative arrival time and absolute arrival time

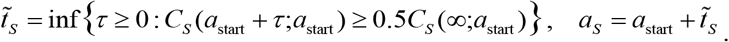

where *a*_start_ denotes the starting age in days of the cohort; 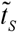 represents the relative arrival time (“reach *S*”); *a*_*S*_ indicates the absolute arrival time; and *C*_*S*_ (∞;*a*_start_ ) represents the total proportion ultimately entering stage *S* . Solutions are obtained via the binary search method (with samples.10^−3^ day precision) and are replicated across bootstrap samples.

### Molecular–morphological alignment

#### Predictive Model for Molecular Ratios

For each molecular marker *m* ∈{*Stra*8, *Sycp*1, *Sycp*3 _ *fil*}, a unified time shift *δ*_*m*_ is introduced along the consolidated meiotic timeline. Its expression profile is synchronized to the same “clock” via a linear combination of the steady-state stage composition *π* _*s*_ (*t*) and the entry flux *J*, with an optional local term *b*_*m*_ :

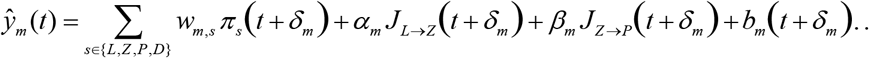

Where *ŷ*_*m*_ (*t*) is the predicted molecular proportion of marker *m* ; *π* _*s*_ (*t*) represents the proportion of the population in stage *s* ; *J*_*L*→*Z*_ (*t*) = *q*_*LZ*_ (*t*)*π* _*L*_(*t*) and *J*_*Z* →*P*_ (*t*) = *q*_*ZP*_ (*t*)*π* _*Z*_ (*t*) represent the entry fluxes given by the piecewise CTMC; *w*_*m,s*_∈[0,1] represents the steady-state stage weights, constrained via nonnegative projection and ridge penalty, approximately satisfying 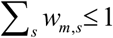 and *α*_*m*_, *β*_*m*_ ≥ 0 denotes the flux weight; *b*_*m*_(·) represents a local term (set to *b*_*Stra*8_^≡ 0^ for Stra8, with the specific forms for Sycp1 and Sycp3_fil described below).

#### Local terms of Sycp1

A narrow Gaussian local term was incorporated to model the early Sycp1 peak on day ∼13.5 −14.5, with concurrent optimization of parameters (*δ, μ,σ, ρ*) :

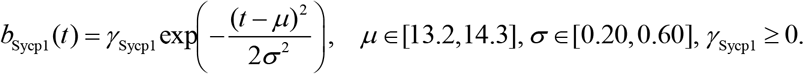

where *μ* denotes the center time (in days) of the pulse; *σ* represents its temporal width (in days); *γ* _Sycp1_ represents the amplitude of this local term; *δ* denotes the global temporal shift (in days); and *ρ* represents the overdispersion of the noise. The specific procedure is as follows: for a set of candidate parameters (*μ,σ,δ* ), perform weighted nonnegative ridge regression to obtain coefficients (*w*_Sycp1,*s*_,*α*_Sycp1_, *β*_Sycp1_,*γ* _Sycp1_ ) . Then, we compute the weighted beta–binomial negative log-likelihood over the candidate values of *ρ*, incorporating a high-weight pseudo-observation that enforces near-zero expression on day 12.5 while assigning an additional weight to data points ≤16 days to prioritize fitting the early minor peak. The parameter combination yielding the lowest score is selected as the final set for Sycp1.

#### Local terms of Sycp3_fil

A mid-stage Gaussian local term was incorporated to model the main *Sycp3_fil* peak at approximately 16–17 days, with concurrent optimization of the parameter *δ* :

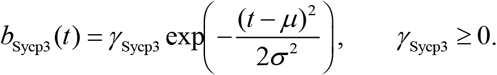

where *μ* denotes the center time (in days) of the pulse; *σ* represents its temporal width (in days); *γ*_Sycp3_ represents the amplitude of this local term; and *δ* denotes the global temporal shift (in days). The specific procedure is as follows: for each candidate value of *δ*, perform weighted nonnegative ridge regression to obtain coefficients (*w*_Sycp1,*s*_,*α*_Sycp1_, *β*_Sycp1_,*γ* _Sycp1_ ) . Then, evaluate the candidates via a weighted sum of squared residuals as the scoring function. This scoring function incorporates a high-weight pseudo-observation enforcing near-zero expression on day 12.5 and assigns a 4× higher weight (e.g., ×4) to data points within the 15–18.5 day window—where the main peak occurs—to ensure that the model prioritizes fitting this mid-stage peak and prevents its dilution by temporal averaging. The candidate *δ* and its corresponding coefficient set yielding the lowest score are selected as the final parameters for Sycp3_fil. Weighted Nonnegative Ridge Regression Estimation

For each molecular marker *m*, after specifying its shape parameters, we apply the same weighted, nonnegativity-constrained ridge regression procedure to solve for the coefficients *θ*_*m*_ :

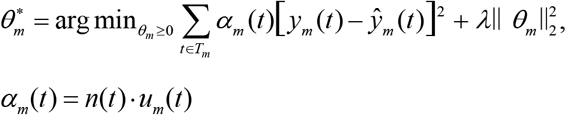

where *λ* = 10^−3^ denotes the ridge penalty coefficient, which serves to stabilize the estimates and prevent excessively large coefficients; *θ*_*m*_={*w*_*m,s*_,*α*_*m*_, *β*_*m*_,*γ* _*m*_} represents the linear combination coefficient vector for this molecular marker *m* ; 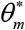 denotes the retained optimal coefficients; *T*_*m*_ represents the set of observation time points for this marker, and *α*_*m*_ (*t*) is the weight assigned to time point *t* ; *y*_*m*_(*t*) denotes the true positive fraction of marker *m* at time point *t* ; *n*(*t*) represents the cell count for population*Vasa*^+^ ; and *u*_*m*_ (*t*) is the corresponding weighting coefficient.

Min–Max normalization

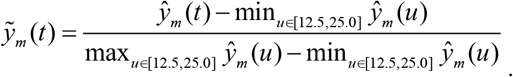

Rise Half-Duration and Full Width at Half Maximum (FWHM)

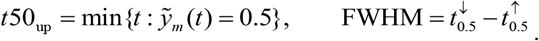

where 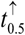 denotes the time of the first occurrence reaching 0.5 and where 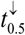 denotes the time of the subsequent return to 0.5; when 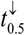 does not exist within the interval [12.5, 25.0], FWHM is recorded as NaN. This study yielded the following results: *Stra*8 *t*50_up_ = 13.48 days, FWHM = NaN ; Sycp1 *t*50_up_ = 13.49 days, FWHM = NaN; and Sycp3_fil *t*50_up_ = 12.983 days, FWHM = 4.805 days.

## Statistical analyses

Statistical analyses were performed using GraphPad Prism 10.6. The quantity of each protein-positive cell number was determined on the basis of a minimum of three independent replicates. Detailed information about the number of independent repetitions, sample size is indicated in the figure legends. The data are presented as the means ± SDs.

## Funding

This work was supported by:

National Natural Science Foundation of China (32030033; 82230052)

The Special Fund for Basic Scientific Research and Business Expenses of Central Universities (63241542)

## Author contributions

Conceptualization: C.L., Z.J., L.L.

Methodology: C.L., Z.J., L.L.

Investigation: C.L., Z.J., G.L., G.F., J.L., Y.W., J.H., L.L.

Visualization: C.L., Z.J., G.L., L.L.

Supervision: C.L., Z.J., J.H., L.L.

Writing—original draft: C.L., Z.J., G.L., L.L.

Writing—review & editing: C.L., Z.J., G.L., L.L.

C.L. and Z.J. designed the experiments, conducted the major experiments, analyzed the data, and prepared the manuscript. G.L. constructed the fitting model. G.F., J.L., and Y.W. conducted some of the experiments and discussed them. L.L. conceived the project, designed the experiments, and revised the manuscript. All the authors read and approved the final paper.

## Competing interests

The authors declare that they have no competing interests.

## Data and materials availability

No unique reagents were generated in this study. GSE128553 from Ge W and GSE136441 from Niu W were downloaded for analysis. All data are available in the main text or the supplementary materials.”

## Supplementary Materials

**Fig. S1.**
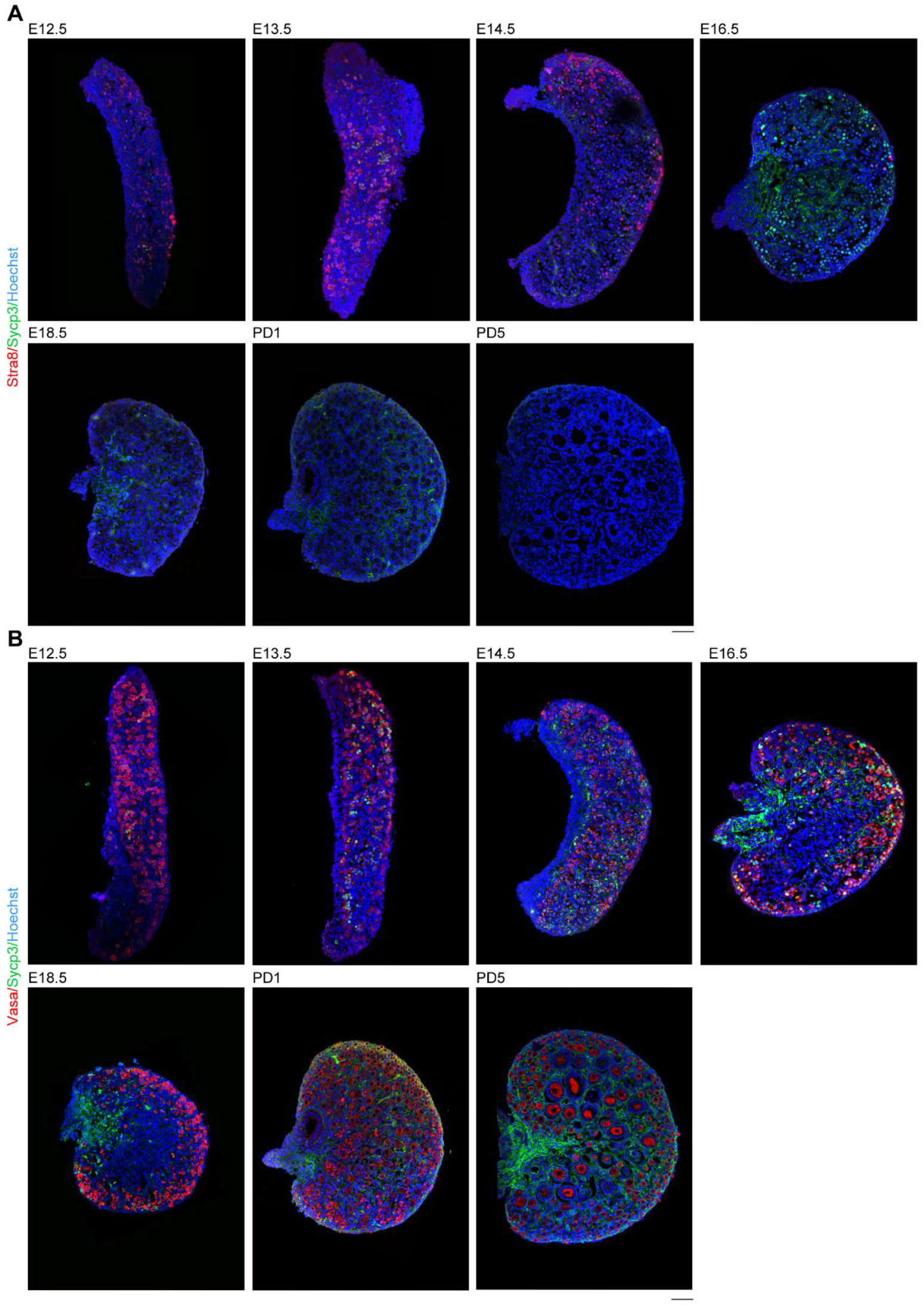
Dynamic expression of Stra8, Vasa and SYCP3 protein in female ovaries. **(A)** Whole co-immunofluorescence images of Stra8 and Sycp3 of *in situ* sections from E12.5, E13.5, E14.5, E16.5, E18.5, PD1 and PD5 embryonic female gonads. Red, Stra8; Green, Sycp3; Blue, nucleus counterstained with Hoechst 33342. Scale bars: 50 μm. **(B)** Whole co-immunofluorescence images of Vasa and Sycp3 of *in situ* sections from E12.5, E13.5, E14.5, E16.5, E18.5, PD1 and PD5 embryonic female gonads. Red, Vasa; Green, Sycp3; Blue, nucleus counterstained with Hoechst 33342. Scale bars: 50 μm. Using Zeiss.LSM710.

**Fig. S2.**
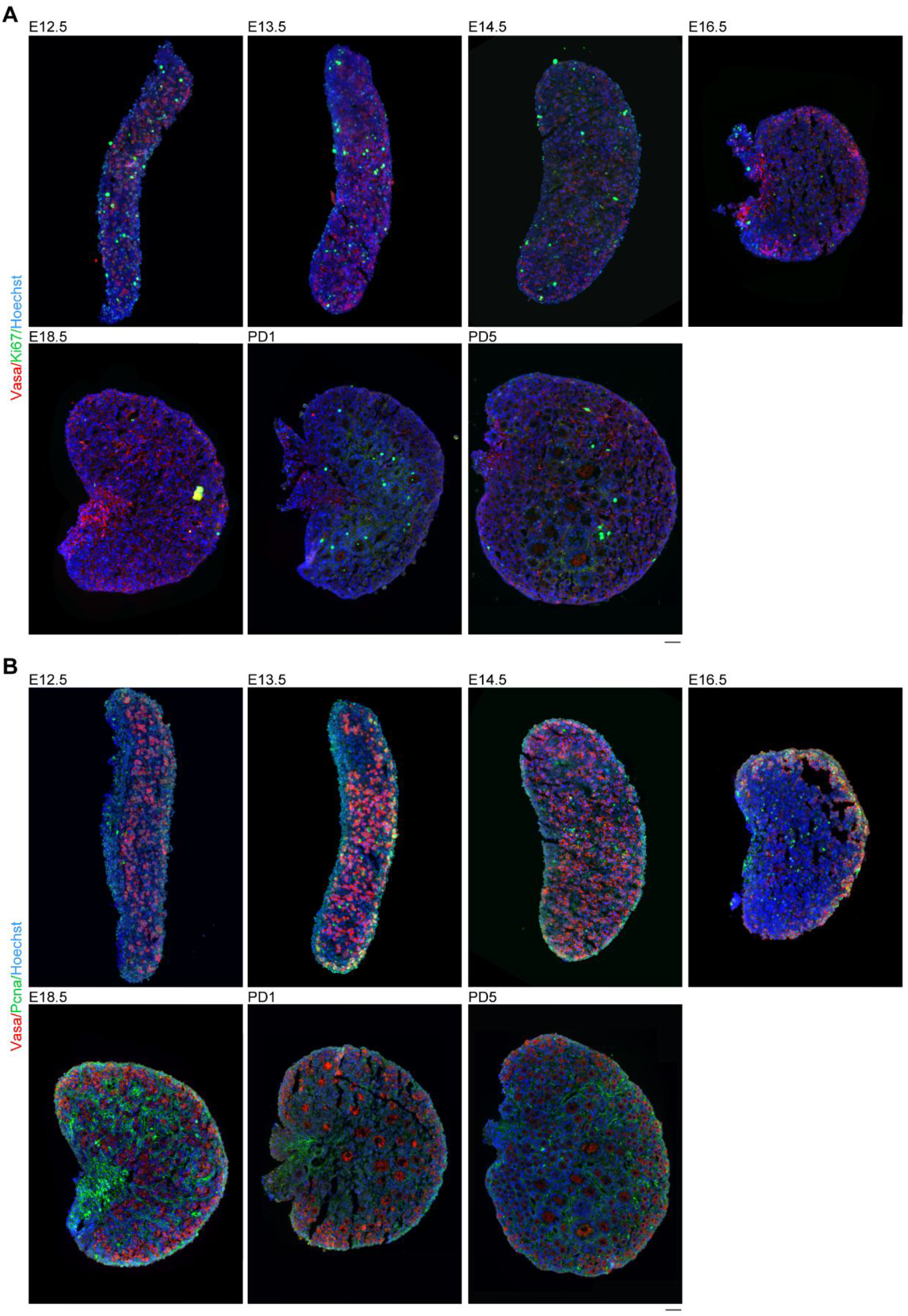
Expression of Ki67, Pcna and Vasa in female ovaries. **(A)** Overall co-immunofluorescence images of Vasa and Ki67 of *in situ* sections from E12.5, E13.5, E14.5, E16.5, E18.5, PD1 and PD5 embryonic female gonads. Red, Vasa; Green, Ki67; Blue, nucleus counterstained with Hoechst 33342. Scale bars: 50 μm. **(B)** Overall co-immunofluorescence images of Vasa and Pcna of *in situ* sections from E12.5, E13.5, E14.5, E16.5, E18.5, PD1 and PD5 embryonic female gonads. Red, Vasa; Green, Pcna; Blue, nucleus counterstained with Hoechst 33342. Scale bars: 50 μm.

**Fig. S3.**
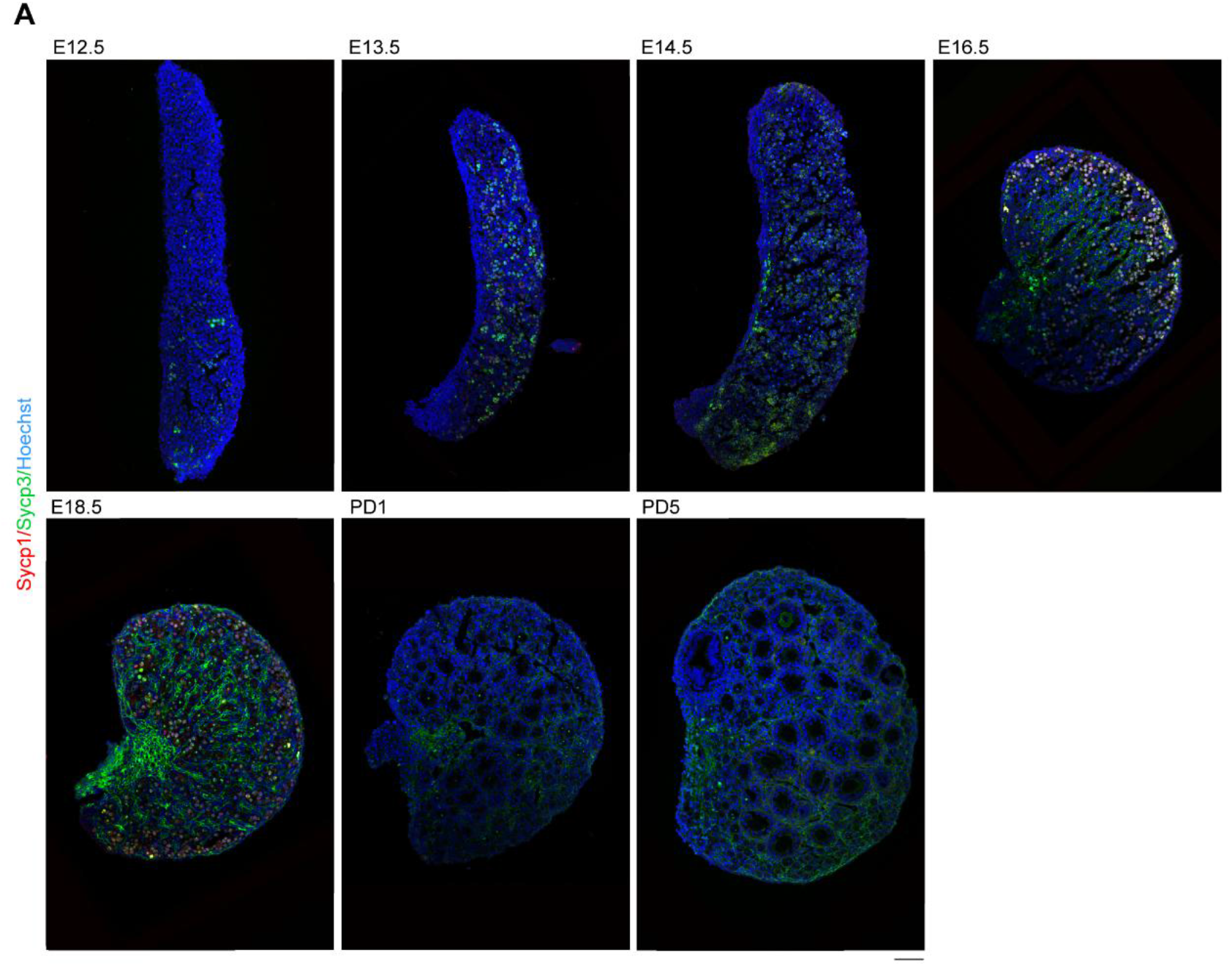
Expression of Ki67, Pcna and Vasa in female ovaries. **(A)** Overall co-immunofluorescence images of Vasa and Ki67 of *in situ* sections from E12.5, E13.5, E14.5, E16.5, E18.5, PD1 and PD5 embryonic female gonads. Red, Vasa; Green, Ki67; Blue, nucleus counterstained with Hoechst 33342. Scale bars: 50 μm. **(B)** Overall co-immunofluorescence images of Vasa and Pcna of *in situ* sections from E12.5, E13.5, E14.5, E16.5, E18.5, PD1 and PD5 embryonic female gonads. Red, Vasa; Green, Pcna; Blue, nucleus counterstained with Hoechst 33342. Scale bars: 50 μm.

**Fig. S4.**
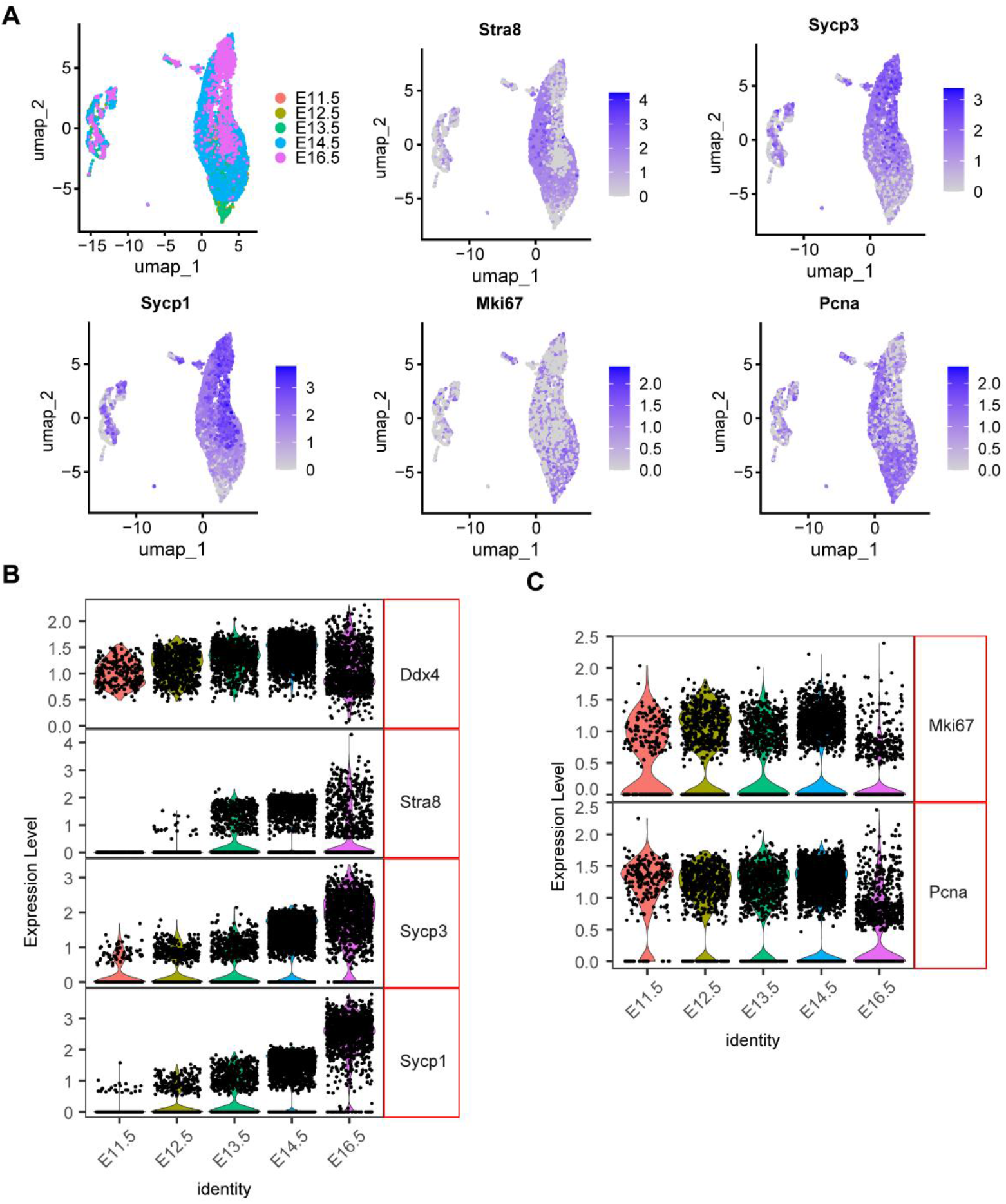
Analysis of 10x sc-RNA-seq data for expression levels of meiosis genes. **(A)** Visualization of clusters using umap (cells are colored by embryonic time points from E11.5 to E16.5). Plots of the expression levels of *Stra8, Sycp3, Sycp1, Ki67* and *Pcna* along the developmental timeline; color intensity indicates the level of expression. **(B)** A multivariate plot showing the relative expression of genes (*Ddx4, Stra8, Sycp3* and *Sycp1*, gene names at right) at different developmental time points (times at the bottom). **(C)** A multivariate plot showing the expression of *Ki67* and *Pcna* at different developmental time points (times at the bottom). All the 10x sc-RNA-seq data used here are from Shen et al.’s article^40^.

**Fig. S5.**
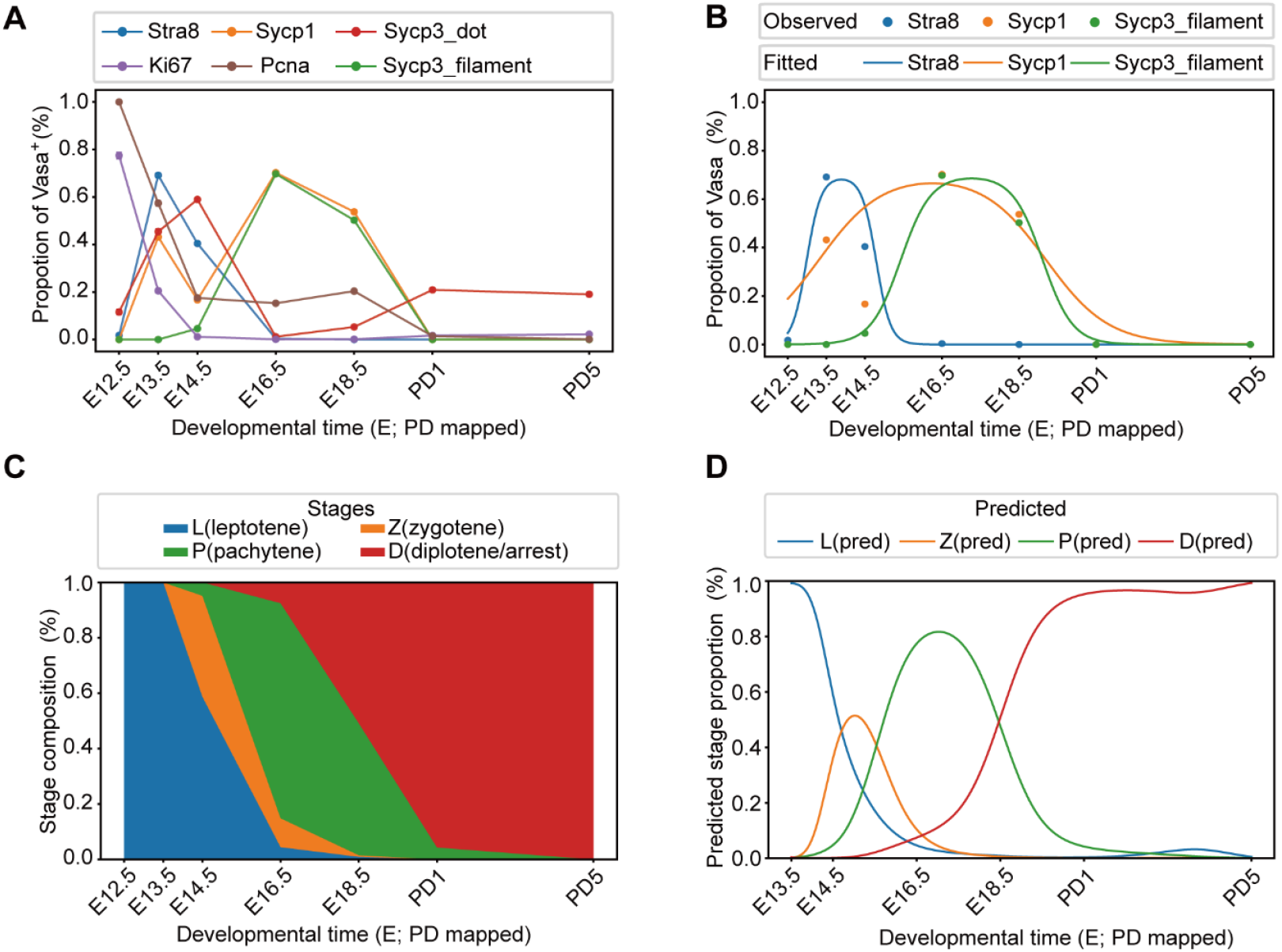
Modeling and Validation of Meiotic Dynamics. **(A)** Observations of marker-positive proportions over development. **(B)** Two-logistic fits of marker-positive proportions over developmental time. **(C)** Stage composition over developmental time: observed (stacked bars). **(D)** Stage composition over time: model predictions (lines).

**Table S1.**
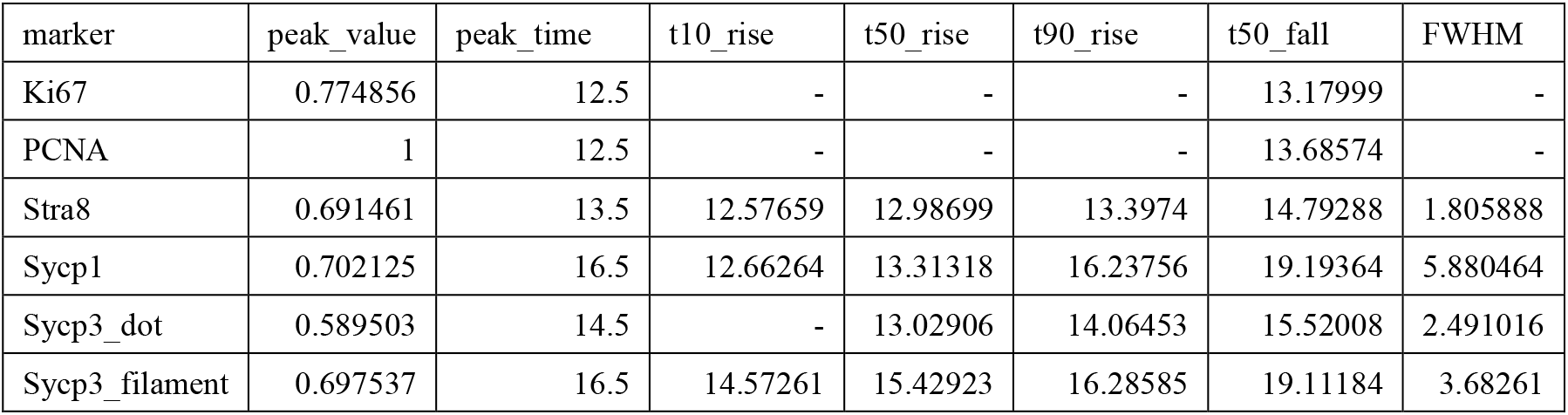
Window metrics ( 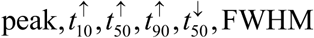) for each marker.

**Table S2.** Per-stage temporal window metrics.

| stage | pmax | tpeak | t10 | t50_up | t90 | t50_down | FWHM | tpeak_label | t50_up_label | t50_down_label | t10_label | t90_label |
| --- | --- | --- | --- | --- | --- | --- | --- | --- | --- | --- | --- | --- |
| L | 0.992 | 13.5 | - | - | - | 14.63837 | - | E13.5 | - | E14.64 | - | - |
| Z | 0.516 | 15.05 | 13.94352 | 14.33498 | 14.71501 | 15.93739 | 1.602 | E15.05 | E14.33 | E15.94 | E13.94 | E14.72 |
| P | 0.818 | 17 | 14.71147 | 15.548 | 16.31731 | 18.65357 | 3.106 | E17.0 | E15.55 | E18.65 | E14.71 | E16.32 |
| D | 0.992 | 24.5 | 16.85997 | 18.46984 | 19.70878 | - | - | PD5.0 | E18.47 | - | E16.86 | PD0.21 |

**Table S3.** Crossing times *Z* > *L, P* > *Z*, and *D* > *P* under the ≥5% rule.

| t_cross_Z_eq<br>uals_L_>=5% | t_cross_P_eq<br>uals_Z_>=5% | t_cross_D_eq<br>uals_P_>=5% | t_cross_Z_equals<br>_L_>=5%_label | t_cross_P_equals<br>_Z_>=5%_label | t_cross_D_equal<br>s_P_>=5%_label |
| --- | --- | --- | --- | --- | --- |
| 14.70048 | 15.54517 | 18.46145 | E14.7 | E15.55 | E18.46 |

**Table S4.** Predicted compositions at selected times.

| time_label | t_numeric | L | Z | P | D |
| --- | --- | --- | --- | --- | --- |
| E13.5 | 13.5 | 0.992015 | 0.001558 | 0.002697 | 0.00373 |
| E13.5 | 13.5 | 0.992015 | 0.001558 | 0.002697 | 0.00373 |
| E14.0 | 14 | 0.917211 | 0.070332 | 0.011836 | 0.00062 |
| E14.5 | 14.5 | 0.587817 | 0.362846 | 0.048172 | 0.001165 |
| E15.0 | 15 | 0.319347 | 0.515596 | 0.159438 | 0.005619 |
| E16.0 | 16 | 0.088884 | 0.235456 | 0.629982 | 0.045678 |
| E16.5 | 16.5 | 0.046298 | 0.103848 | 0.775352 | 0.074503 |
| E17.0 | 17 | 0.026821 | 0.044284 | 0.817793 | 0.111102 |
| E18.0 | 18 | 0.012542 | 0.010852 | 0.671173 | 0.305433 |
| E18.5 | 18.5 | 0.008825 | 0.005867 | 0.475656 | 0.509652 |
| PD1.0 | 20.5 | 0.004063 | 0.000835 | 0.042851 | 0.952251 |
| PD5.0 | 24.5 | 0.006154 | 0.000651 | 0.00092 | 0.992275 |

**Table S5.**
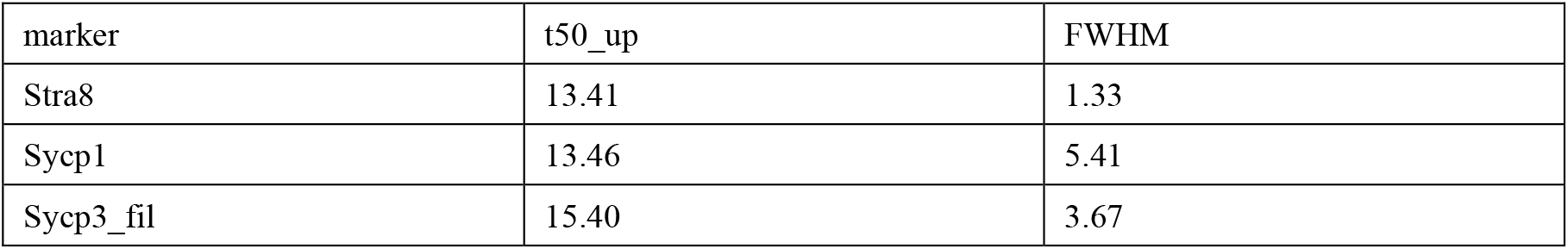
Molecular Proportion Window Metrics: Rise Half-Duration *t*50_up_ and FWHM (Min–Max Normalized)

## Notes

### Competing Interest Statement

The authors have declared no competing interest.

## REFERENCES

1. M. E. Pepling, A. C. Spradling, Female mouse germ cells form synchronously dividing cysts. Development 125, 3323–3328 (1998).

2. K. J. Grive, R. N. Freiman, The developmental origins of the mammalian ovarian reserve. Development 142, 2554–2563 (2015).

3. H. Larose, A. N. Shami, H. Abbott, G. Manske, L. Lei, S. S. Hammoud, “Gametogenesis: A journey from inception to conception” in Organ Development, D. M. Wellik, Ed. (Current Topics in Developmental Biology, 2019), vol. 132, pp. 257–310.

4. J. Bowles, P. Koopman, Retinoic acid, meiosis and germ cell fate in mammals. Development 134, 3401–3411 (2007).

5. L. M. Mehlmann, Stops and starts in mammalian oocytes: recent advances in understanding the regulation of meiotic arrest and oocyte maturation. Reproduction 130, 791–799 (2005).

6. J. Bowles, D. Knight, C. Smith, D. Wilhelm, J. Richman, S. Mamiya, K. Yashiro, K. Chawengsaksophak, M. J. Wilson, J. Rossant, H. Hamada, P. Koopman, Retinoid signaling determines germ cell fate in mice. Science 312, 596–600 (2006).

7. X. Y. Wang, M. E. Pepling, Regulation of Meiotic Prophase One in Mammalian Oocytes. Front. Cell. Dev. Biol. 9, 12 (2021).

8. K. Borum, Oogenesis in the mouse--A study of the meiotic prophase. Experimental cell research 24, 495–507 (1961).

9. L. D. Jonathan M. W. Slack, “Germ cell and gonadal development” in Essential Developmental Biology, 4th Edition (John Wiley & Sons Ltd, 2022), pp. 329–330.

10. D. B. Menke, J. Koubova, D. C. Page, Sexual differentiation of germ cells in XX mouse gonads occurs in an anterior-to-posterior wave. Dev. Biol. 262, 303–312 (2003).

11. M. Bullejos, P. Koopman, Germ cells enter meiosis in a rostro-caudal wave during development of the mouse ovary. Mol. Reprod. Dev. 68, 422–428 (2004).

12. R. Shimada, Y. Kato, N. Takeda, S. Fujimura, K. Yasunaga, S. Usuki, H. Niwa, K. Araki, K. Ishiguro, STRA8-RB interaction is required for timely entry of meiosis in mouse female germ cells. Nat. Commun. 14, 18 (2023).

13. A. E. Baltus, D. B. Menke, Y. C. Hu, M. L. Goodheart, A. E. Carpenter, D. G. de Rooij, D. C. Page, In germ cells of mouse embryonic ovaries, the decision to enter meiosis precedes premeiotic DNA replication. Nature Genetics 38, 1430–1434 (2006).

14. L. Yuan, J. G. Liu, J. Zhao, E. Brundell, B. Daneholt, C. Höög, The murine “SCP3” gene is required for synaptonemal complex assembly, chromosome synapsis, and male fertility. Molecular Cell 5, 73–83 (2000).

15. J. Koubova, D. B. Menke, Q. Zhou, B. Capel, M. D. Griswold, D. C. Page, Retinoic acid regulates sex-specific timing of meiotic initiation in mice. Proc. Natl. Acad. Sci. U. S. A. 103, 2474–2479 (2006).

16. E. L. Anderson, A. E. Baltus, H. L. Roepers-Gajadien, T. J. Hassold, D. G. de Rooij, A. M. M. van Pelt, D. C. Page, “Stra8” and its inducer, retinoic acid, regulate meiotic initiation in both spermatogenesis and oogenesis in mice. Proc. Natl. Acad. Sci. U. S. A. 105, 14976–14980 (2008).

17. M. D. Griswold, C. A. Hogarth, J. Bowles, P. Koopman, Initiating Meiosis: The Case for Retinoic Acid. Biol. Reprod. 86, 7 (2012).

18. D. Zickler, N. Kleckner, Recombination, Pairing, and Synapsis of Homologs during Meiosis. Cold Spring Harbor Perspect. Biol. 7, 26 (2015).

19. L. Yuan, J. G. Liu, J. Zhao, E. Brundell, B. Daneholt, C. Höög, The murine SCP3 gene is required for synaptonemal complex assembly, chromosome synapsis, and male fertility. Molecular Cell 5, 73–83 (2000).

20. J. Fraune, S. Schramm, M. Alsheimer, R. Benavente, The mammalian synaptonemal complex: Protein components, assembly and role in meiotic recombination. Experimental Cell Research 318, 1340–1346 (2012).

21. L. Yuan, J. G. Liu, M. R. Hoja, J. Wilbertz, K. Nordqvist, C. Höög, Female germ cell aneuploidy and embryo death in mice lacking the meiosis-specific protein SCP3. Science 296, 1115–1118 (2002).

22. K. K. Billmyre, E. A. Kesler, D. Tsuchiya, T. J. Corbin, K. Weaver, A. Moran, Z. L. Yu, L. Adams, K. Delventhal, M. Durnin, O. R. Davies, R. S. Hawley, SYCP1 head-to-head assembly is required for chromosome synapsis in mouse meiosis. Science Advances 9, (2023).

23. H. Wang, Q. C. Liu, S. F. Cheng, L. Li, W. Shen, W. Ge, Single-Cell Transcriptomic Analysis of the Potential Mechanisms of Follicular Development in Stra8-Deficient Mice. International Journal of Molecular Sciences 26, (2025).

24. G. A. Dokshin, A. E. Baltus, J. J. Eppig, D. C. Page, Oocyte differentiation is genetically dissociable from meiosis in mice. Nature Genetics 45, 877–U289 (2013).

25. S. M. Guo, Y. R. Zhang, C. F. Fei, X. Z. Liu, W. Xia, M. C. Luo, G. H. Wei, W. B. Qin, C. L. Xiong, H. G. Li, Y. Yin, X. M. He, L. Q. Zhou, Deciphering meiotic chromatin organization by SYCP3. Nucleic Acids Research 53, (2025).

26. J. H. Lammers, H. H. Offenberg, M. van Aalderen, A. C. Vink, A. J. Dietrich, C. Heyting, The gene encoding a major component of the lateral elements of synaptonemal complexes of the rat is related to X-linked lymphocyte-regulated genes. Molecular and cellular biology 14, 1137–1146 (1994).

27. D. H. Castrillon, B. J. Quade, T. Y. Wang, C. Quigley, C. P. Crum, The human VASA gene is specifically expressed in the germ cell lineage. Proc. Natl. Acad. Sci. U. S. A. 97, 9585–9590 (2000).

28. Y. Fujiwara, T. Komiya, H. Kawabata, M. Sato, H. Fujimoto, M. Furusawa, T. Noce, Isolation of a DEAD-family protein gene that encodes a murine homolog of Drosophila vasa and its specific expression in germ cell lineage. Proc. Natl. Acad. Sci. U. S. A. 91, 12258–12262 (1994).

29. Y. Toyooka, N. Tsunekawa, Y. Takahashi, Y. Matsui, M. Satoh, T. Noce, Expression and intracellular localization of mouse Vasa-homologue protein during germ cell development. Mechanisms of Development 93, 139–149 (2000).

30. K. Vaithiyanathan, S. H. Liew, N. Zerafa, T. Gamage, M. Cook, L. A. O’Reilly, P. Bouillet, C. L. Scott, A. Strasser, J. K. Findlay, K. J. Hutt, BCL2-modifying factor promotes germ cell loss during murine oogenesis. Reproduction 151, 553–562 (2016).

31. M. Myers, F. H. Morgan, S. H. Liew, N. Zerafa, T. U. Gamage, M. Sarraj, M. Cook, I. Kapic, A. Sutherland, C. L. Scott, A. Strasser, J. K. Findlay, J. B. Kerr, K. J. Hutt, PUMA regulates germ cell loss and primordial follicle endowment in mice. Reproduction 148, 211–219 (2014).

32. J. B. Kerr, R. Duckett, M. Myers, K. L. Britt, T. Mladenovska, J. K. Findlay, Quantification of healthy follicles in the neonatal and adult mouse ovary: evidence for maintenance of primordial follicle supply. Reproduction 132, 95–109 (2006).

33. M. Balla, R. Angelopoulou, G. Lavranos, C. Kittas, Follicular cells versus oocytes: Cell population dynamics in the developing ovary. Tissue & Cell 40, 373–381 (2008).

34. R. G. Lea, L. P. Andrade, M. T. Rae, L. T. Hannah, C. E. Kyle, J. F. Murray, S. M. Rhind, D. W. Miller, Effects of maternal undernutrition during early pregnancy on apoptosis regulators in the ovine fetal ovary. Reproduction 131, 113–124 (2006).

35. M. Biot, A. Toth, C. Brun, L. Guichard, B. de Massy, C. Grey, Principles of chromosome organizationfor meiotic recombination. Molecular Cell 84, 1826–1841 (2024).

36. G. Hamer, I. Novak, A. Kouznetsova, C. Höög, Disruption of pairing and synapsis of chromosomes causes stage-specific apoptosis of male meiotic cells. Theriogenology 69, 333–339 (2008).

37. K. A. Ferguson, V. Chow, S. Ma, Silencing of unpaired meiotic chromosomes and altered recombination patterns in an azoospermic carrier of a t(8;13) reciprocal translocation. Human Reproduction 23, 988–995 (2008).

38. A. Di Carlo, G. Travia, M. De Felici, The meiotic specific synaptonemal complex protein SCP3 is expressed by female and male primordial germ cells of the mouse embryo. Int. J. Dev. Biol. 44, 241–244 (2000).

39. W. B. Niu, A. C. Spradling, Two distinct pathways of pregranulosa cell differentiation support follicle formation in the mouse ovary. Proc. Natl. Acad. Sci. U. S. A. 117, 20015–20026 (2020).

40. W. Ge, J. J. Wang, R. Q. Zhang, S. J. Tan, F. L. Zhang, W. X. Liu, L. Li, X. F. Sun, S. F. Cheng, P. W. Dyce, M. De Felici, W. Shen, Dissecting the initiation of female meiosis in the mouse at single-cell resolution. Cellular and Molecular Life Sciences 78, 695–713 (2021).

41. Z. L. Pei, K. Deng, C. J. Xu, S. Zhang, The molecular regulatory mechanisms of meiotic arrest and resumption in Oocyte development and maturation. Reprod. Biol. Endocrinol. 21, 12 (2023).

42. M. Saitou, H. Miyauchi, Gametogenesis from Pluripotent Stem Cells. Cell Stem Cell 18, 721–735 (2016).

43. J. He, A. Yan, B. Chen, J. H. Huang, K. Kee, 3D genome remodeling and homologous pairing during meiotic prophase of mouse oogenesis and spermatogenesis. Dev. Cell 58, 3009–3027 (2023).

44. M. A. Handel, J. C. Schimenti, Genetics of mammalian meiosis: regulation, dynamics and impact on fertility. Nature Reviews Genetics 11, 124–136 (2010).

45. B. Chen, G. Z. Zhu, A. Yan, J. P. He, Y. Liu, L. P. Li, X. R. Yang, C. P. Dong, K. Kee, IGSF11 is required for pericentric heterochromatin dissociation during meiotic diplotene. Plos Genetics 17, (2021).

46. S. Jiang, Z. Huang, Y. Li, C. W. Yu, H. Yu, Y. W. Ke, L. Jiang, J. Liu, Single-cell chromatin accessibility and transcriptome atlas of mouse embryos. Cell Reports 42, (2023).

47. M. Y. Liu, Y. Yin, X. Y. Ye, M. Zeng, Q. Zhao, D. L. Keefe, L. Liu, Resveratrol protects against age-associated infertility in mice. Human Reproduction 28, 707–717 (2013).

48. J. E. Holt, V. Pye, E. Boon, J. L. Stewart, I. García-Higuera, S. Moreno, R. Rodríguez, K. T. Jones, E. A. McLaughlin, The APC/C activator FZR1 is essential for meiotic prophase I in mice. Development 141, 1354–U1327 (2014).

49. K.-i. Ishiguro, K. Matsuura, N. Tani, N. Takeda, S. Usuki, M. Yamane, M. Sugimoto, S. Fujimura, M. Hosokawa, S. Chuma, M. S. H. Ko, K. Araki, H. Niwa, MEIOSIN Directs the Switch from Mitosis to Meiosis in Mammalian Germ Cells. Dev. Cell 52, 429–445 (2020).

50. L. Liu, S. Franco, B. Spyropoulos, P. B. Moens, M. A. Blasco, D. L. Keefe, Irregular telomeres impair meiotic synapsis and recombination in mice. Proc. Natl. Acad. Sci. U. S. A. 101, 6496–6501 (2004).

51. S. X. Fan, Y. W. Wang, H. W. Jiang, X. H. Jiang, J. T. Zhou, Y. Y. Jiao, J. W. Ye, Z. S. Xu, Y. Wang, X. F. Xie, H. Zhang, Y. Li, W. Liu, X. J. Zhang, H. Ma, B. L. Shi, Y. W. Zhang, M. Zubair, W. S. Shah, Z. P. Xu, B. Xu, Q. H. Shi, A novel recombination protein C12ORF40/REDIC1 is required for meiotic crossover formation. Cell Discovery 9, (2023).

52. Y. H. Hao, T. Stuart, M. H. Kowalski, S. Choudhary, P. Hoffman, A. Hartman, A. Srivastava, G. Molla, S. Madad, C. Fernandez-Granda, R. Satija, Dictionary learning for integrative, multimodal and scalable single-cell analysis. Nature Biotechnology 42, 293–304 (2024).

